# Identification of a Distinct Metabolomic Subtype of Sporadic ALS Patients

**DOI:** 10.1101/416396

**Authors:** Qiuying Chen, Davinder Sandhu, Csaba Konrad, Dipa Roychoudhury, Benjamin I. Schwartz, Roger R. Cheng, Kirsten Bredvik, Hibiki Kawamata, Elizabeth L. Calder, Lorenz Studer, Steven. M. Fischer, Giovanni Manfredi, Steven. S. Gross

## Abstract

Sporadic amyotrophic lateral sclerosis (sALS) is a progressive motor neuron disease resulting in paralysis and death. Genes responsible for familial ALS have been identified, however the molecular basis for sALS is unknown. To discover metabotypic biomarkers that inform on disease etiology, untargeted metabolite profiling was performed on 77 patient-derived dermal fibroblast lines and 45 age/sex-matched controls. Surprisingly, 25% of sALS lines showed upregulated methionine-derived homocysteine, channeled to cysteine and glutathione (GSH). Stable isotope tracing of [U-^13^C]-glucose showed activation of the trans-sulfuration pathway, associated with accelerated glucose flux into the TCA cycle, glutamate, GSH, alanine, aspartate, acylcarnitines and nucleotide phosphates. A four-molecule support vector machine model distinguished the sALS subtype from controls with 97.5% accuracy. Plasma metabolite profiling identified increased taurine as a hallmark metabolite for this sALS subset, suggesting systemic perturbation of cysteine metabolism. Furthermore, integrated multiomics (mRNAs/microRNAs/metabolites) identified the super-trans-sulfuration pathway as a top hit for the sALS subtype. We conclude that sALS can be stratified into distinct metabotypes, providing for future development of personalized therapies that offer new hope to sufferers.

## Introduction

Amyotrophic lateral sclerosis (ALS) is a fatal neurodegenerative disease characterized by progressive death of upper and lower motor neurons; symptoms include muscle weakness and atrophy, leading to fatal paralysis, usually within 5 years of disease onset^1^. Approximately 90% of ALS patients have no familial history (a condition termed sporadic ALS; sALS), while the remaining 10% are due to recognized inherited gene mutations^2^. Unfortunately, mechanisms leading to motor neuronal death in sALS are completely unknown and, as a consequence, there are neither viable disease biomarkers nor effective therapies. ALS clinical trials have been largely unsuccessful^3^, and currently there are only two approved drugs, Riluzole and Edaravone, both of which only prolong the lifetime of ALS cases by a few months. Conceivably, sALS can be driven by multiple mechanistic triggers, and disease stratification (i.e., recognition of distinct disease-driver subtypes) could lead to effective targeted/personalized therapeutic approaches.

As the apparent heterogeneity of the sALS patient population presents a major stumbling stone to rigorous mechanistic studies, stratifying sALS has become an area of considerable interest^4^, ^5^ The worldwide *ALS Stratification Prize - Using the Power of Big Data and Crowdsourcing for Catalyzing Breakthroughs in ALS* was initiated 2015, aiming to address the problem of ALS patient heterogeneity with consideration of important clinical distinctions, such as changes in the ALS functional rating scale^6^. Stratification efforts have identified several potential non-standard predictors of disease progression, including plasma uric acid, creatinine and surprisingly, blood pressure, perhaps related to sALS pathobiology^6^. While this stratification relies on clinical parameters to anticipate disease progression, recognition of disease-associated metabolic perturbations are likely to be more telling, with potential for stratification of sALS patients based on metabotypes that identify distinct molecular disease mechanisms that potentially inform rationally-targeted new therapies.

It has been shown that altered metabolism, including increased resting energy expenditure, often precedes the clinical onset of ALS^7^. Moreover, alterations in energy metabolism have been inferred as a potential pathogenic mechanism underlying sALS^8^, ^9^ Notably, as the final downstream products of genes, transcripts and proteins, metabolites offer the most proximal insights into sALS disease mechanisms. Whereas specific metabolic perturbations that underlie bioenergetic defects in sALS have remained elusive, understanding this knowledge is likely to inform on disease mechanisms that prove transformational for clinical care. Accordingly, we recognize metabolites as the ultimate readout of the genome /trancriptome/proteome, with potential to stratify sALS cases and inform personalized therapies.

Here, we sought to investigate whether the metabotype of skin-derived fibroblasts from sALS patients can enable patient subtyping and disease stratification. Notably, fibroblasts carry the same genetic composition as neurons, the relevant tissue in ALS disease, but unlike neural tissue, fibroblasts can be readily accessible from cases for *in vitro* growth and analysis. Indeed, prior studies of dermal fibroblast metabolism have been employed to reveal bioenergetic alterations in sALS cases^5, 9, 10^. Here, we performed the first untargeted metabolite profiling on patient-derived dermal fibroblasts, comparing relative metabolite abundances with age- and sex-matched fibroblasts from healthy control subjects. This investigation revealed a unique subset of sALS case fibroblasts that display a distinct shared metabotype, typified by accelerated trans-sulfuration pathway-derived cysteine for support of GSH biosynthesis, along with hypermetabolism of glucose. The finding that sALS patients can be classified based on metabotype offers a potentially critical first-step towards novel targeted personalized medicines and effective new disease therapies.

## Methods

### Reagents

LC-MS grade acetonitrile (ACN), isopropanol (IPA) and methanol (MeOH) were purchased from Fischer Scientific. High purity deionized water (ddH2O) was filtered from Millipore (18 OMG). OmniTrace glacial acetic acid and ammonium hydroxide were obtained from EMD Chemicals. [U-13C] glucose, [2,3,3-^2^H]- serine were purchased from Cambridge Isotope Laboratory. Ammonium acetate and all other chemicals and standards were obtained from Sigma Aldrich in the best available grade.

### Cell culture

A total of 77 sALS and 45 control fibroblast cell lines were obtained from cases and propogate as previously described (Konrad et al. Mol Neuorodeg 2017). 75,000 cells/well for each line were plated in duplicate 6-well plates and cultured in DMEM medium containing 5 mM glucose and 4 mM glutamine, 10% FBS,1% of 100X antibiotic/antimycotic (Ab+F; which contains sterile-filtered 10,000 units penicillin, 10 mg streptomycin and 25 μg amphotericin B per mL, and 2.5 µg/ml Plasmocin). Cells were harvested at 80% confluency and extracted for LC/MS metabolomic analysis.

### Metabolite extraction

Each cell line was cultured and initially extracted as two biological replicates, for independent LC/MS metabolomic data acquisition. Cells were washed twice with ice-cold PBS, followed by metabolite extraction using −70°C 80:20 methanol:water (LC-MS grade methanol, Fisher Scientific). The tissue–methanol mixture was subjected to bead-beating for 45 sec using a Tissuelyser cell disrupter (Qiagen). Extracts were centrifuged for 5 min at 5,000 rpm to pellet insoluble material and supernatants were transferred to clean tubes. The extraction procedure was repeated two additional times and all three supernatants were pooled, dried in a Vacufuge (Eppendorf) and stored at −80°C until analysis. The methanol-insoluble protein pellet was solubilized in 0.2 M NaOH at 95°C for 20 min and protein was quantified using a BioRad DC assay. On the day of metabolite analysis, dried cell extracts were reconstituted in 70% acetonitrile at a relative protein concentration of 1 µg/ml, and 4 µl of this reconstituted extract was injected for LC/MS-based untargeted metabolite profiling.

These 244 patient-derived skin fibroblast extracts (122 samples with 2 biological replicates) were analyzed by LC-QTOF metabolomics profiling in random sequence. To adjust for day-to-day and batch-to-batch LC/MS instrument drift, metabolite stability, and other experimental factors that may contribute to systematic error, metabolite measurements were normalized to one another using flanking quality control (QC) samples run at intervals of every 6 injections. These QC samples were prepared from a pool of all samples and this normalization procedure enabled the comparative analysis of data acquired over a 1-month collection period.

Plasma metabolites were extracted by addition to 1 part plasma to 20 parts 70% acetonitrile in ddH2O (vol:vol). The mixture was briefly vortexed and then centrifuged for 5 min at 16,000 × g to pellet precipitated proteins. An aliquot of the resulting extract (3 μl) was subjected to untargeted metabolite profiling using, applying both positive and negative ion monitoring MS.

### Untargeted metabolite profiling by LC/MS

Cell extracts were analyzed by LC/MS as described previously^11,12,^ using a platform comprised of an Agilent Model 1290 Infinity II liquid chromatography system coupled to an Agilent 6550 iFunnel time-of-flight MS analyzer. Chromatography of metabolites utilized aqueous normal phase (ANP) chromatography on a Diamond Hydride column (Microsolv). Mobile phases consisted of: (A) 50% isopropanol, containing 0.025% acetic acid, and (B) 90% acetonitrile containing 5 mM ammonium acetate. To eliminate the interference of metal ions on chromatographic peak integrity and electrospray ionization, EDTA was added to the mobile phase at a final concentration of 6 µM. The following gradient was applied: 0-1.0 min, 99% B; 1.0-15.0 min, to 20% B; 15.0 to 29.0, 0% B; 29.1 to 37min, 99% B. Raw data were analyzed using MassHunter Profinder 8.0 and MassProfiler Professional (MPP) 14.9.1 software (Agilent technologies). Mann Whitney t-tests (p<0.05) were performed to identify significant differences between groups.

### Metabolite Structure Specification

To ascertain the identities of differentially expressed metabolites (P<0.05), LC/MS data was searched against an in-house annotated personal metabolite database created using MassHunter PCDL manager 7.0 (Agilent Technologies), based on monoisotopic neutral masses (<5 ppm mass accuracy) and chromatographic retention times. A molecular formula generator (MFG) algorithm in MPP was used to generate and score empirical molecular formulae, based on a weighted consideration of monoisotopic mass accuracy, isotope abundance ratios, and spacing between isotope peaks. A tentative compound ID was assigned when PCDL database and MFG scores concurred for a given candidate molecule. Tentatively assigned molecules were verified based on a match of LC retention times and/or MS/MS fragmentation spectra for pure molecule standards contained in a growing in-house metabolite database.

### Stable isotope tracing of [U-^13^C] glucose and [2,3,3-^2^H] serine

An in-house untargeted stable isotope tracing (USIT) workflow was employed using Agilent metabolite profiling software MassHunter Qualitative Analysis 7.0, MassProfinder 8.0 and MPP 14.9.1. Labelled metabolites were identified on the basis of differential abundance in cells cultured with supplemental heavy isotope-labelled metabolite vs. natural (light) isotope, as described previously^13,14^ Selected metabolites were identified on the basis of previously curated isotopologues. Notably, the USIT workflow calculates and corrects for the natural abundance of ^13^C and ^2^H isotope in samples.

### Microarray analysis

Total RNA (combined mRNA and miRNA) from skin fibroblasts was extracted from 27 sALS patients and 27 controls using Trizol agent (Invitrogen). Total RNA was further extracted using the Agilent Absolutely RNA miRNA Kit. The total RNA concentration and RNA sample integrity were verified by a Bio-Analyzer 2100 (Agilent Technologies, Waldbronn, Germany). The quality of isolated RNA was determined using an Agilent 2200 TapeStation system and Bioanalyzer with mRNA labeling and microarray processing were performed according to the manufacturers recommendations. miRNA labeling was done using an Agilent miRNA Complete Labeling and Hyb Kit with gene expression and miRNA data extracted using Agilent Feature Extraction Software.

Extracted mRNA and miRNA data were analyzed using the respective workflows in GeneSpring GX 14.9.1 (Agilent Technologies, CA). Signal intensities for each mRNA probe was normalized to 75th percentile values with baseline transformation for gene expression analysis. Separately, miRNA data was normalized to 90th percentile and baseline transformed for miRNA analysis. The differential miRNA list (P<0.05, FC>1.2) was used to identify the gene targets based on a target prediction database incorporated in GeneSpring GX (e.g. TargetScan, PicTar, microRNA.org). The differentially expression mRNA was combined with validated miRNA targets and metabolites for integrated multi-Omics pathway analysis using databases linked to KEGG, Wiki pathways and Biocyc.

### Statistics

All values are averages of at least three independent measurements. Error bars indicate standard deviation (S.D.) or standard error of the mean (S.E.M.). Statistically significant differences between two groups were estimated by unpaired two-tailed Student’s test with significance set at p<0.05.

## Results

### Untargeted metabolite profiling identifies a distinct sALS case metabotype

Table 1 summarizes the clinical characteristics of 77 sALS cases and 45 healthy controls studied herein. Dermal fibroblast cell lines were established from all cases and 1172 quality control-normalized metabolite features were detected with at least 70% of the resulting cell cultures. Metabolite profiling and non-parametric Mann-Whitney statistical analysis (P<0.05) revealed that 507 of these 1172 recognized features were differentially expressed in sALS fibroblasts. We next performed unsupervised machine learning algorithm principal component analysis (PCA) to determine if the heterogeneous sALS cohort could be confidently distinguished from the control population. Whereas sALS as a whole did not completely separate from the control group (Fig 1B), a distinct subgroup of 18/77 sALS case-derived fibroblast clustered together in this PCA analysis, separating from the controls based on principal component 1 (PC1), which accounted for 33.9% of total metabolite variance.

**Table 1.** Clinical characteristics of the sALS and control cohorts

| **Subject** | **sALS-Male** | **sALS-Female** | **sALS-Total** | **CTR-Male** | **CTR-Female** | **CTR-Total** |
| --- | --- | --- | --- | --- | --- | --- |
| **N** | 43 | 34 | 77 | 23 | 22 | 45 |
| **BMI** | 26.54 ± 4.55 | 26.04 ± 6.41 | 26.32 ± 5.41 | NA | NA | NA |
| **Age at Symptom Onset** | 56.28 ± 9.92 | 62.44 ± 8.45 | 59.00 ± 9.74 | NA | NA | NA |
| **Age at Time of Skin BX** | 57.65 ± 9.76 | 63.858 ± 0.44 | 60.39 ± 9.66 | 64.83 ± 8.95 | 57.32 ± 5.40 | 61.15 ± 8.27 |
| **ALSFRS-R Total at Biopsy** | 32.81 ± 8.61 | 32.38 ± 7.80 | 32.62 ± 8.21 | NA | NA | NA |
| **% Rate of Decline** | 1.20 ± 0.84 | 1.08 ± 0.56 | 1.141 ± 0.72 | NA | NA | NA |
| **FVC (%)** | 74.78 ± 25.61 | 75.84 ± 29.05 | 75.25 ± 27.00 | NA | NA | NA |

Specified values are means ± S.D. ALSFRS-R = ALS functional rating scale–revised. FVC = Forced vital capacity.

**Figure 1.**
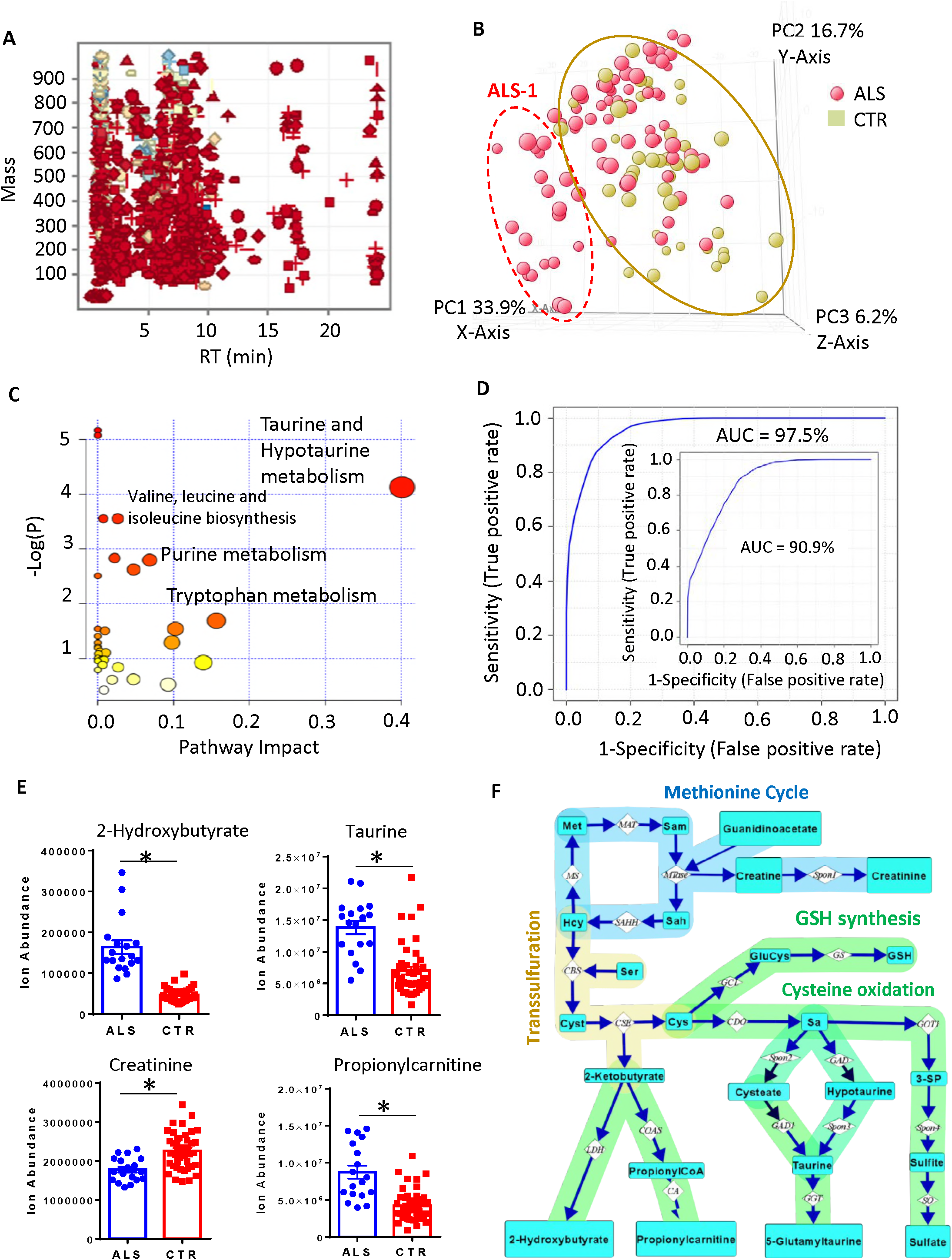
Untargeted metabolite profiling of case-derived fibroblast lines identifies an sALS subgroup with an enhanced trans-sulfuration pathway and glucose hypermetabolism. Panel A: mass and retention time (RT) scatter plot of 1165 features detected at 70% frequency in at least one group of 77 ALS and 45 controls. Panel B; PCA score plot showing incomplete separation of 77 sALS (red circles) from 45 controls (yellow circles), but a subgroup of >18 sALS patients (identified in the dashed red oval as sALS-1) was readily separated from all controls. Panel C: integrated enrichment and pathway topology analysis of differential metabolites in ALS-1 by Metaboanalyst 3.0, recognizing taurine and hypotaurine in the trans-sulfuration pathway as the most significant pathway impact and lowest matching P-value. Panel D: *ROC* curve derived from SVM mode predicting >18 sALS subgroup from a group of 45 randomly selected control fibroblasts. The AUC (i.e. the overall accuracy, specificity and sensitivity) of the prediction is 97.5%. The inset showed the *ROC* curve and AUC for a repeat experiment comprising the same 18 sALS subset vs. 18 randomly selected control fibroblast cell lines. Panel E: Ion abundance of 4 sALS-1 predictor metabolites (2-hydroxybutyrate, taurine, propionylcarnitine and creatinine) from 18 sALS-1 and 45 controls; *, P<0.05 (2-tailed Student t-test). Panel F: Schematic representation of metabolites and enzymes involved in the methionine cycle, methylation and trans-sulfuration pathways. Abbreviation of metabolite names and enzymes are as follows: Met, methionine; Hcy: homocystieine; Sam: S-adenosylmethionine; Sah: S-adenosylhomocysteine; Cyst: cystathionine; Sa: sulfoalanine; 3-SP: 3-sulfopyruvate. The enzyme name abbreviations were adapted from KEGG pathway. MAT: methionine adenosyltransferase; MS: methionine synthase; SAHH: S-adenosylhomocysteine hydrolase; MTase: methyltransferase; CBS: cstathionine beta synthase; CSE: cystathionine gamma lyase; GCL: glutamate cysteine ligase; GS: GSH synthase; CDO: cysteine dioxygenase; LDH: lactate dehydrogenase; COAS: propionylCoA synthetase; CA: carnitine acyltransferase; GAD1: glutamate decarboxylase 1; GGT: Gamma-glutamyltransferase; GOT1: glutamic-oxaloacetic transaminase 1; SO: sulfite oxidase; Spon1, Spon2 and Spon3 refer to non-enzymatic spontaneous degradation process.

To confirm that the sALS subgroup (denoted sALS-1) could be reproducibly distinguished from the control group and other sALS patients (denoted sALS-2), we repeated the metabolic profiling in an independent repeat analysis, comparing the same 18 sALS-1 patient subgroup vs. a group of 18 control fibroblast lines randomly selected from the original 45 control lines. Results confirmed a group of 41 metabolites, including both structurally-identified and structurally–unidentified species, to be differentially-expressed in both the original study (18 sALS vs. 45 control cell lines) and the confirmation study (18 sALS vs. 18 controls) (Fig S1). Included in the group of 41 differentially-expressed metabolites, significant increases were observed for 2-hydroxybutyrate, taurine, and C3-C5 carnitines, along with significant decreases in creatinine, ATP and several amino acids. Integrated enrichment and pathway topology analysis of these differentially-expressed metabolites, performed using Metaboanalyst 3.1, revealed the taurine and hypotaurine pathway to be the most significantly impacted (Fig 1C).

Using a support vector machine (SVM) learning algorithm and ROC-curve based biomarker analysis, we generated a 4-metabolite linear SVM model (comprising 2-hydroxybutyrate, taurine, propionylcarnitine and creatinine) that predicted the sALS subgroup with an overall accuracy of 97% in the original group (with 1/45 false-positives and 1/18 false-negatives; Fig 1D) and 90.9% accuracy in the repeat analysis. Notably, targeted quantitative analysis reconfirmed the veracity of the observed decreased level of creatinine and increased levels of 2-hydroxybutyrate, taurine, and propionylcarnitine (Fig 1E).

Remarkably, this group of 4 metabolites share interconnectivity in the methionine cycle, methylation and trans-sulfuration pathways (Fig 1F). Notably, under metabolic stress conditions, supplies of L-cysteine for glutathione synthesis can become limiting, causing the normal regeneration of methionine from homocysteine, to diversion of homocysteine into the trans-sulfuration pathway for production of cystathionine as a first step in cysteine biosynthesis. Cystathionine is cleaved to cysteine and 2-ketobutyrate, and the latter can be enzymatically reduced to 2-hydroxybutyrate. Alternatively, 2-ketobutyrate can enter the TCA cycle via addition to propionylcarnitine for fatty acid β-oxidation in mitochondria (Fig 1F). Creatine is synthesized from guanidinoacetate in a S-adenosylmethionine (SAM) dependent reaction^15^ and can spontaneously degrade to creatinine (Fig 1F). It has been shown that up to 40% of SAM is consumed for methylation of creatine^16^. Diminished intracellular creatinine suggests a diminished rate of SAM-mediated methylation in skin fibroblasts from sALS-1 cases. Collectively, these untargeted metabolite profiling findings indicate significant trans-sulfuration/methylation pathway perturbations in fibroblasts isolated from sALS-1 patients.

### Serine incorporation into glutathione is increased in the trans-sulfuration enhanced sALS metabotype

We next asked whether the observed increases in trans-sulfuration pathway metabolites may be linked to an increased demand for glutathione synthesis in the sALS-1 subgroup. In one approach to assess this possibility, ALS-1 and control fibroblasts were grown in 2 mM [2,3,3-^2^H]-serine supplemented medium, followed by a determination of the extent of deuterium enrichment in GSH. Three sALS fibroblasts with highest intracellular 2-hydroxyglutarate were chosen for this stable isotope tracing experiment, for comparison with 3 randomly selected control fibroblast cell lines. Notably, serine metabolism to glycine would predictably result in a singly-deuterated glycine that may be incorporated into *de novo* synthesized GSH. Indeed, fibroblast growth in medium supplemented with [2,3,3-^2^H]-serine demonstrated a modest, but significantly increased isotopic incorporation into GSH from sALS cell lines vs. controls at 1h, 5h, and 24h, (Fig 2A, B). Similarly, GSSG showed a significantly elevated serine-derived glycine incorporation in the sALS-1 group fibroblasts, along with a significant increase in absolute GSSG levels vs. control at 5h and 24h (Fig 2C, D).

**Figure 2.**
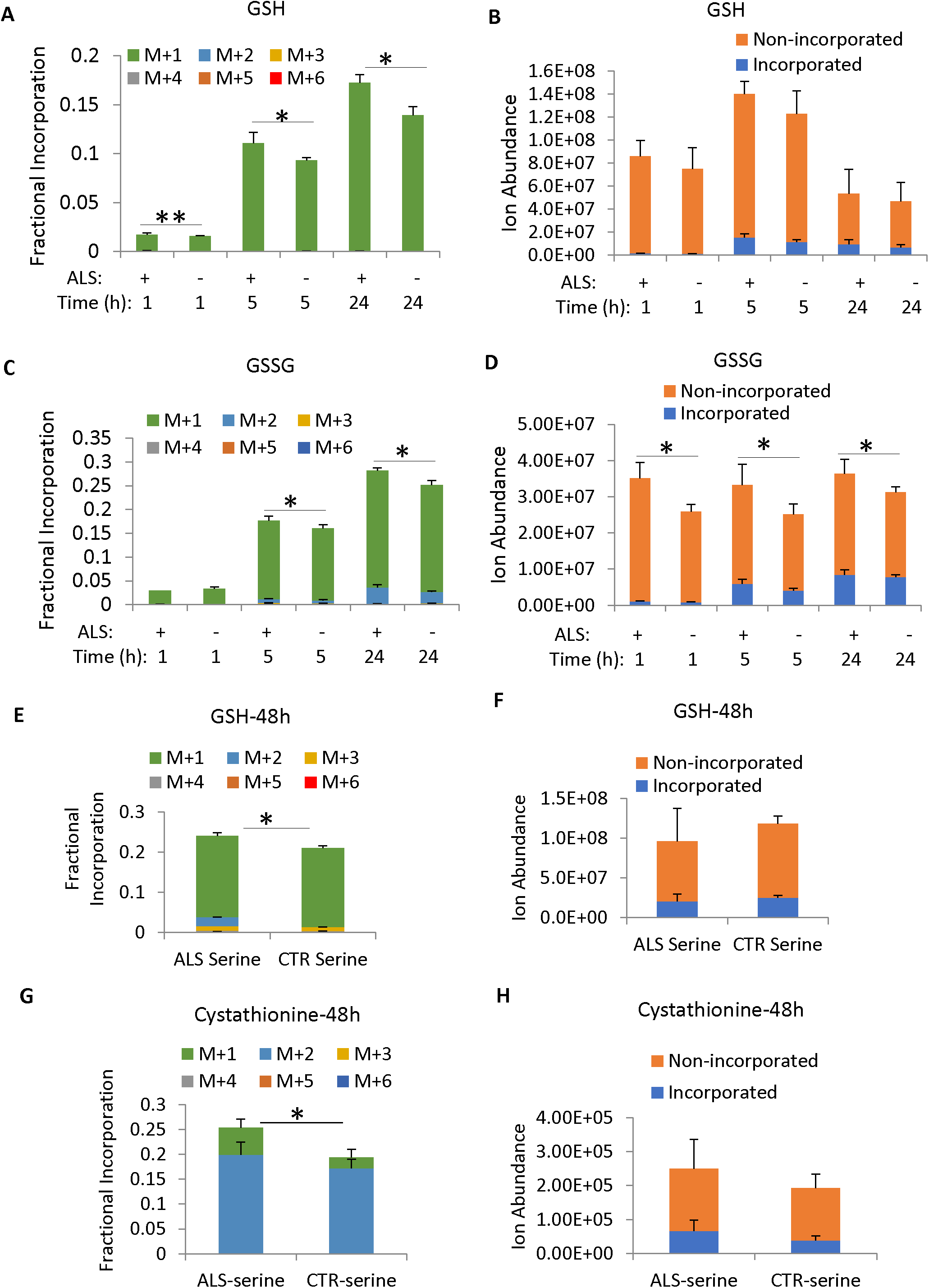
Increased incorporation of [2,3,3-^2^H]-serine into GSH, GSSG and cystathionine synthesis in the sALS-1 group after 1h, 5h, 24h and 48h in culture. Panel A and B: fractional isotopic enrichment (A) and total ion abundance (B) of GSH after 1h, 5h, and 24h culture in [2,3,3-^2^H]-serine isotope enrichment. Panel C and D:isotopic enrichment (C) and ion abundance (D) of GSSG after 1h, 5h, and 24h of culture in [2,3,3-^2^H]-serine. Panel E and F: fractional isotopic enrichment (E) and ion abundance (F) of GSH after 48h culture in of [2,3,3-^2^H]-serine. Panel G-H: fractional isotopic enrichment (G) and ion abundance (H) of cystathionine after 48h of culture in [2,3,3-^2^H]-serine. Data was obtained from 3 independent sALS-1 vs. 3 control cell lines and presented as mean ± S.D. *, P<0.05 (2-tailed Student t-test).

The biosynthesis of cytosolic GSH is known to be tightly regulated by cysteine availability, and the activity of γ-glutamylcysteinyl ligase (GCL) ^17^-^19^. As shown in Fig 1F, cystathionine is synthesized by cystathionine β-synthase (CBS) via condensation of homocysteine with serine. Hence, the extent of deuterium-labeled serine incorporated into cystathionine can serve to assess activity of the trans-sulfuration pathway. Surprisingly, we did not detect any serine incorporation into cystathionine at 1h, 5h and 24h after [2,3,3-^2^H]- serine supplementation, indicating that trans-sulfuration from homocysteine to *de novo* synthesized cysteine was insignificant during this period. One explanation for this apparent lack of de novo cysteine production is that cystine in the culture medium was ample to support GSH production over a 24h period in culture

To assess whether activation of trans-sulfuration is indeed opposed by cysteine/cystine availability in the culture medium, we cultured cells in [2,3,3-^2^H]-serine supplemented medium for a longer duration of 48h, a time when cystine is significantly decreased. At this time, increased incorporation of serine-derived cysteine for GSH synthesis was indeed observed in the sALS-1 group vs. control (as demonstrated by M+2 and M+3 isotopologues, as well as a predominant serine-derived M+1 glycine incorporation), while the total GSH pool was not significantly different (Fig 2E, F). In accord with utilization of serine for *de novo* cysteine synthesis from cystathionine, after 48 h we also observed incorporation of deuterium from serine into cystathionine, and increased incorporation into the sALS-1 subgroup vs. control (Fig 2G-H). The relatively small fraction of serine-derived M+2 and M+3 isotopologues in GSH suggests that a larger portion of cystathionine-derived cysteine was directed toward oxidation to taurine and reduction to hydrogen sulfide, as schematized in Fig 1F. Therefore, increased cysteine demand in the sALS subgroup, at least in part for GSH synthesis, resulted in accelerated trans-sulfuration of homocysteine to cysteine, concomitant with increased NADH dependent reduction of alpha-ketobutyrate to 2-hydroxybutyrate (Fig 1F). It is notable that the cystathionine gamma-lyase/hydrogen sulfide system was previously reported to be essential for maintaining cellular glutathione status^20^. Accordingly, upregulation of the trans-sulfuration pathway in the sALS-1 patient subgroup can serve to maintain levels of GSH, providing a potential mechanism for cell protection against oxidant production and oxidative stress.

### [U-^13^C]-glucose incorporation into GSH synthesis is increased in the sALS trans-sulfuration metabotype

In addition to serine-derived glycine and cysteine, we explored the relative incorporation of glutamate into GSH synthesis. Glucose, the dominant carbon source for most cells, enters the TCA cycle as acetyl-CoA and contributes carbon atoms to produce α-ketoglutarate and its transamination product, glutamate. We compared the indirect contribution of [U-^13^C]-glucose-derived glutamate to GSH synthesis in fibroblast cell lines; 3 sALS-1 vs. 3 controls as employed for the [2,3,3-^2^H]-serine tracing experiment in Fig 2. To this end, 5 mM [U-^13^C]-glucose was added to glucose-free DMEM medium and ^13^C incorporation into glutamate was quantified after cell growth at 1h, 5h, 24h and 48h. As early as 1h after supplementation, ^13^C enrichment of intracellular glucose approximated 100%, and incorporation into glutamate was observed at all time points predominantly as the M+2 isotopologue (Fig 3A-B). As expected, sALS fibroblasts also showed significantly increased GSH incorporation from glucose-derived glutamate (M+2) after growth in [U-^13^C]-glucose containing medium at 24h and 48h (Fig 3C, E). Notably, [U-^13^C]-glucose can also incorporate into serine, and serine-derived glycine can be further incorporated into GSH. However, under the cell culture conditions used, incorporation of glucose into glycine (M+1) was much less than glutamate M+2 (Fig 3C-E). Despite increases in GSH incorporation from glutamate at 24h, the total GSH pool remained unchanged in sALS vs. control groups (Fig 3D, F), suggesting increased GSH consumption in the sALS-1 group. This suggestion is supported by the observation of increased incorporation ^13^C in GSSG and an increase in the total GSSG pool in sALS-1 vs, control fibroblasts (Fig 3G-H). This finding indicates accelerated GSH oxidation to GSSG and enhanced levels of NADPH consumption as a means to provide for enhanced demand for antioxidant defense in the sALS-1 group.

**Figure 3.**
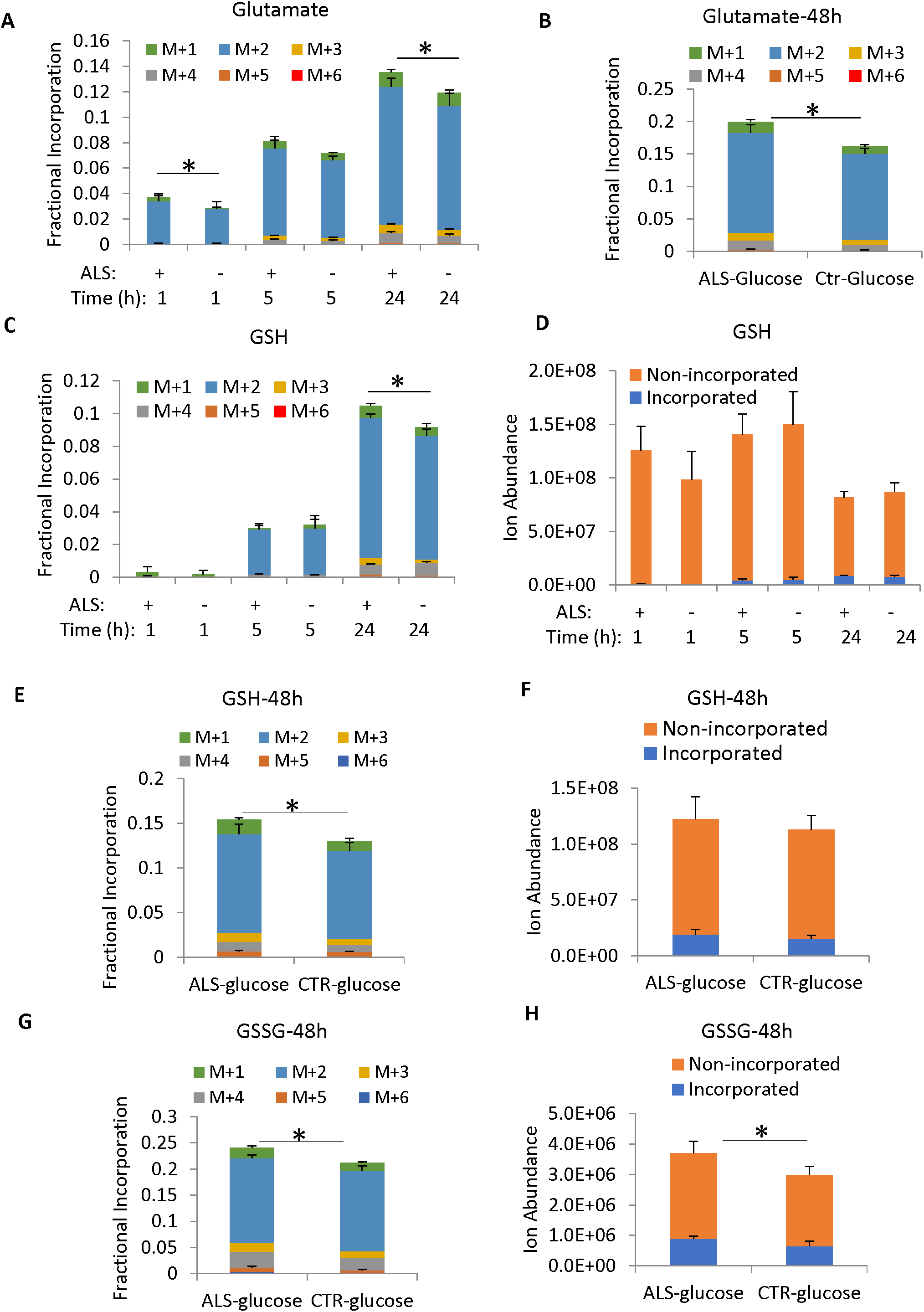
Glutamate, GSH and GSSG fractional incorporation and ion abundance after 1h, 5h, 24h and 48h of[U-^13^C] glucose isotope enrichment. Data was obtained from 3 independent sALS-1 vs. 3 control cell lines and presented as mean ± S.D. *, P<0.05 (2-tailed Student t-test).

### sALS subgroup fibroblasts exhibit accelerated [U-^*13*^C]-glucose incorporation into TCA cycle intermediates and additional pathway metabolites

Stable isotope tracing studies also shed light on whether increased GSH synthesis was associated with an accelerated overall metabolic rate, considering glycolysis, the TCA cycle, and other linked metabolic pathways. Toward this end, we analyzed targeted and untargeted ^13^C tracing results from cells grown in [U-^13^C]-glucose medium for 1h, 5h and 24h. Notably, sALS fibroblasts exhibited a significantly increased fractional incorporation of [U-^13^C]-glucose into lactate, TCA cycle intermediates, amino acids (aspartate and alanine), and C2 and C4 acyl-carnitines (Fig 4A-F and Fig 5A-D). Interestingly, regardless of various changes in metabolite abundance (Fig S2-3), the relative incorporation of glucose into each of these products increased at 24h. Glycolytic intermediates, phosphoenolpyruvate, 2-phosphoglycerate, glucose 6-phosphate and fructose 6-phosphate, similarly showed a trend toward increased glucose incorporation in the sALS-1 group compared to controls after 24h in culture, whereas the relative changes were much smaller compared to TCA cycle intermediates and other pathways (Fig S4). However, we observed increased TCA cycle and cataplerotic products in an independent experiment, where cells were cultured for 48h in [U-^13^C]-glucose-supplemented medium (Fig S5). Interestingly, most TCA cycle intermediates exhibited <15% enrichment from ^13^C in glucose, except for citrate which exhibited a 40% incorporation after 24h. This is consistent with the fact that rapidly proliferating fibroblasts operate a truncated TCA cycle where instead of oxidation to oxoglutarate, citrate is diverted to cytosol for oxidation to oxoglutarate by isocitrate dehydrogenase 1 (IDH1) concomitant with NADPH production and generation of cytosolic acetyl-CoA for fatty acid synthesis^21^. Our data indicate that the sALS fibroblast subgroup exhibit an accelerated TCA cycle flux, both oxidative and reductive, a feature supportive of increased energy metabolism. This apparent hypermetabolic phenotype is in accord with that which we previously described based on analysis of mitochondrial bioenergetics in sALS fibroblasts^5, 9^

**Figure 4.**
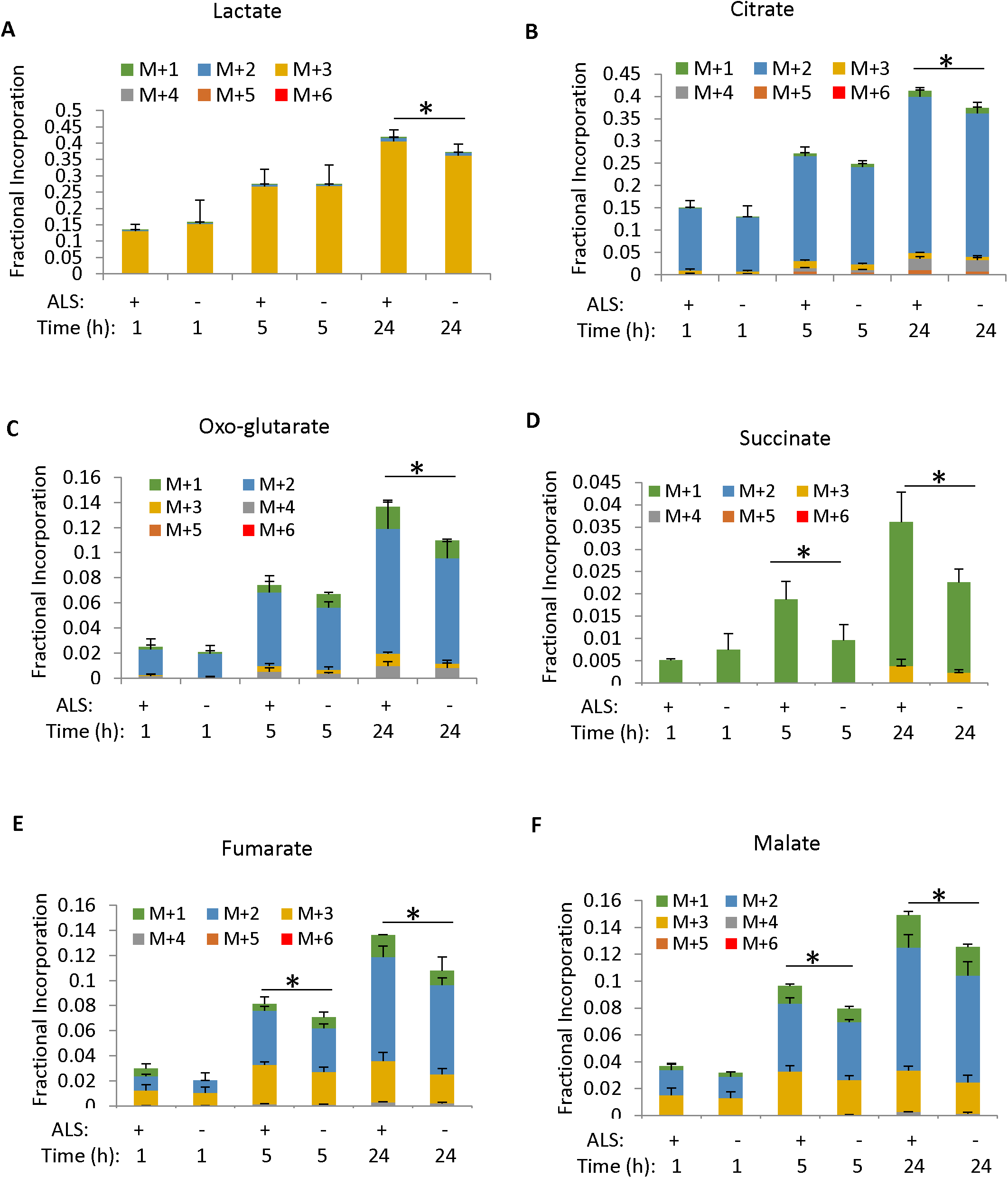
Fractional incorporation of [U-^13^C] glucose into lactate and TCA cycle intermediates after 1h, 5h and 24h of isotope enrichment. Note: Due to the overlapping of succinate M+2 with the reference ion (m/z 119.03) used for mass accuracy correction, the calculated M+2 incorporation in succinate was much smaller than the actual incorporated value. However, the relative incorporation of [U-^13^C] glucose to succinate is significantly increased in sALS-1 compared to the control under the same condition. Data was obtained from 3 independent sALS-1 vs. 3 control cell lines and presented as mean ± S.D. *, P<0.05 (2-tailed Student t-test).

**Figure 5.**
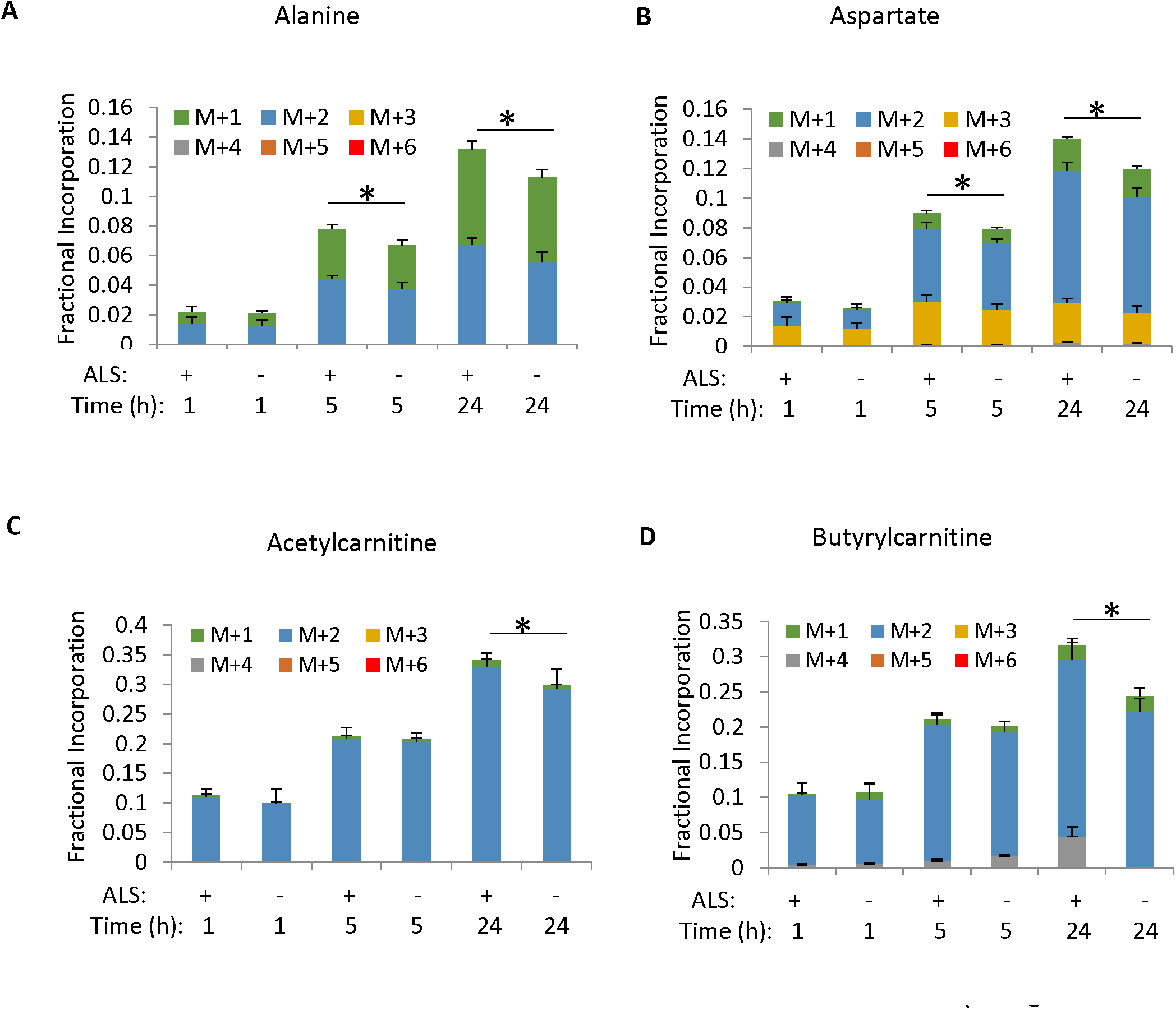
Fractional incorporation of [U-^13^C] glucose into amino acid and acylcarnitines after 1h, 5h and 24h of isotope enrichment. Data was obtained from 3 independent sALS-1 vs. 3 control cell lines and presented as mean ± S.D. *, P<0.05 (2-tailed Student t-test).

In accord with an accelerated TCA cycle, sALS fibroblasts revealed significantly increased glucose incorporation into nucleotide triphosphate pools after 24 h (mostly via the ribose M+5 isotopologue), including ATP, GTP, UTP and CTP, while the individual nucleotide triphosphate total abundances remained unchanged (Fig 6, A-D, Fig S6). This can be explained by sALS-1 fibroblasts adopting an increased rate of both *de novo* nucleotide synthesis and consumption of purine/pyrimidine nucleotide phosphates, thereby maintaining relatively unchanged total intracellular nucleotide pools. Notably, 6-phosphogluconate, a precursor for synthesis of NADPH, ribulose-5-phosphate and ribose 5-phosphate production via the pentose phosphate pathway, showed increased fractional enrichment in sALS (Fig 6E). Increased incorporation of glucose into the oxidative phase of pentose phosphate pathway occurred rapidly, reaching 85% by 1h. Taken together, these findings reveal that sALS-1 fibroblasts exhibit a faster utilization of glucose for nucleotide synthesis and NADPH production, associated with increased glucose metabolism and *de novo* GSH generation.

**Figure 6.**
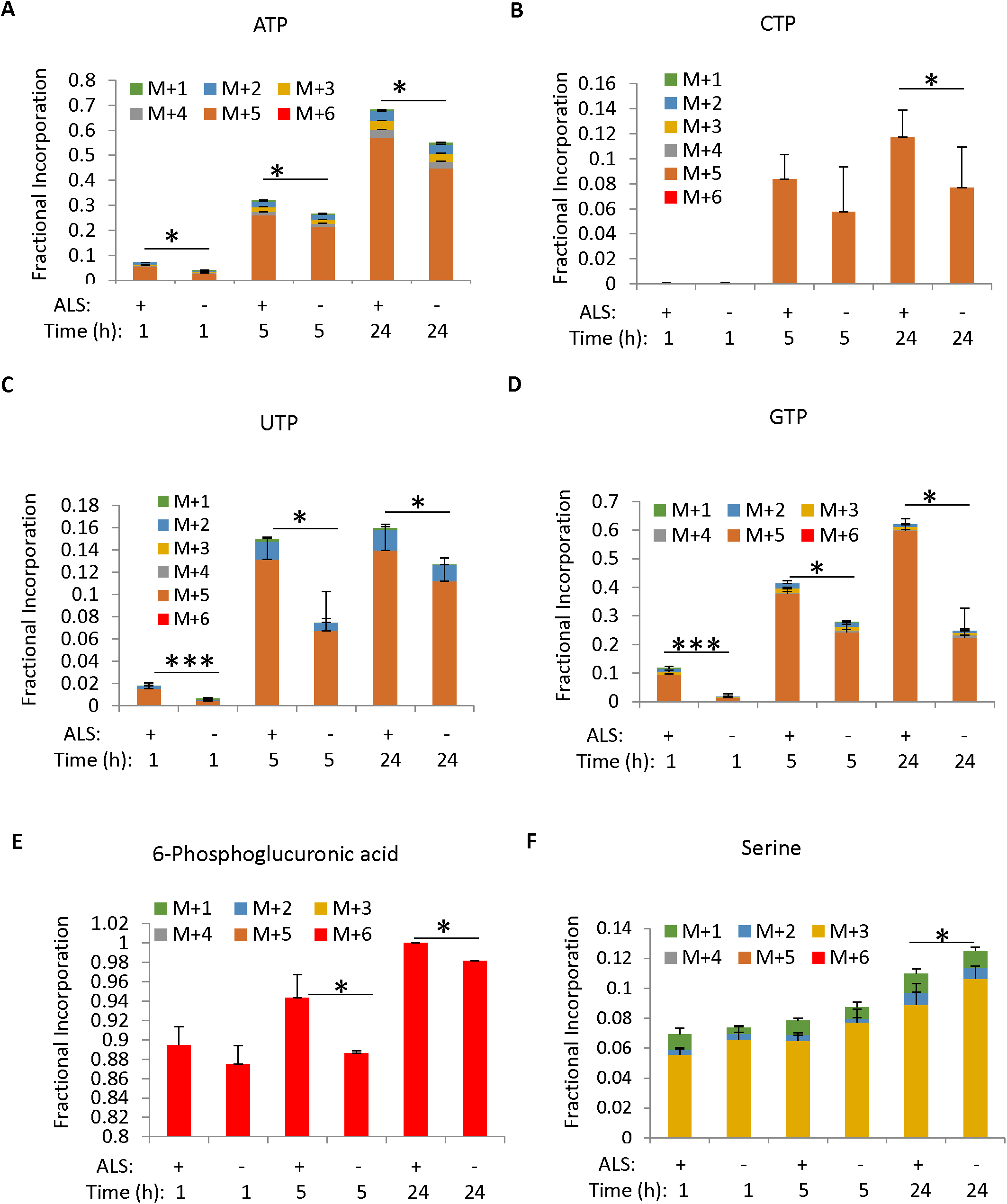
Fractional incorporation of [U-^13^C] glucose into nucleotide triphosphates, 6-phosphoglucuronate and serine synthesis after 1h, 5h and 24h of isotope enrichment. Data was obtained from 3 independent sALS-1 vs. 3 control cell lines and presented as mean ± S.D. *, P<0.05 (2-tailed Student t-test).

Although glucose incorporation into purine nucleotide phosphates was increased, serine synthesis from glucose-derived phosphoglycerate was relatively decreased in sALS-1 fibroblasts (Fig 6F). Serine is the major one-carbon source in proliferating cells for *de novo* synthesis of purine nucleotides. Decreased glucose shunting to serine biosynthesis might be attributed to the fact this study was performed using non-dialyzed serum. Under these conditions, purine salvage from serum provides the dominant carbon source for nucleotide biosynthesis (unpublished finding). Because we did not observe any difference in the rate of sALS-1 cell proliferation compared to control fibroblasts, relative differences in purine metabolism cannot be explained by growth rate differences.

Taken together, our results indicate that sALS fibroblasts exhibit an increased glucose influx to the TCA cycle, cataplerotic enrichment of amino acid synthesis, accelerated GSH synthesis, and accelerated pentose phosphate pathway activity, providing antioxidant NADPH and ribose for nucleotide synthesis.

### Integrated transcriptomic and metabolomic pathway analysis

We performed microarray and microRNA analysis to investigate whether altered glucose and trans-sulfuration metabolism can be explained by selective transcription changes in sALS-1 case fibroblasts, relative to control fibroblasts and sALS-2 fibroblasts that did not conform to the transsulfuration/hyper-metabolism metabotype of sALS-1 cells. Toward this end, transcripts and microRNA targets in fibroblasts from a total of 27 sALS cases compared to transcripts in fibroblasts from 27 randomly selected control subjects. Of the 27 sALS patient cell lines studied, 11 belonged to the sALS-1 metabotype and 16 were sALS-2. Due to the comprehensive nature of gene annotation databases (mostly cancer and disease related), we first curated the searchable database to include potential candidate gene sets based on previously published sALS studies, then identified targeted genes based on differentially-expressed transcripts and microRNA targets in sALS subgroups. Finally, transcriptomic data were overlaid on metabolite profiling data for multiomic data integration and interpretation.

Using the curated database, Gene Set Enrichment analysis (GSA) identified significantly differences in the expression of genes involved in the regulation of catabolic processes and apoptosis in sALS-1 fibroblasts (Fig S7). Interestingly, gene sets related to these two biological processes were significantly upregulated in sALS-1 relative to sALS-2 and control groups. Additionally, epigenetic regulation by H3K4Me2 appeared to be activated in sALS-1 compared to sALS-2 and control. Epigenetic regulation of senescence showed no difference between sALS-1 and sALS-2 and was suppressed in both sALS groups compared to control. Based on GSA findings, we integrated transcriptome, microRNAome and metabolome data into metabolic pathways queried by KEGG, Wiki-pathways and BioCyc. The results indicated that super-trans-sulfuration pathway was among the top list of pathways significantly enriched in sALS-1 (P<2.3e-4 by Fischer’s exact test, Fig 7A). Note that sALS-1 fibroblasts showed a much smaller P-value enrichment than sALS-2 cells, in accord with the trans-sulfuration pathway being significantly accelerated in sALS-1 fibroblasts vs. the sALS-2 and control group cells.

**Figure 7.**
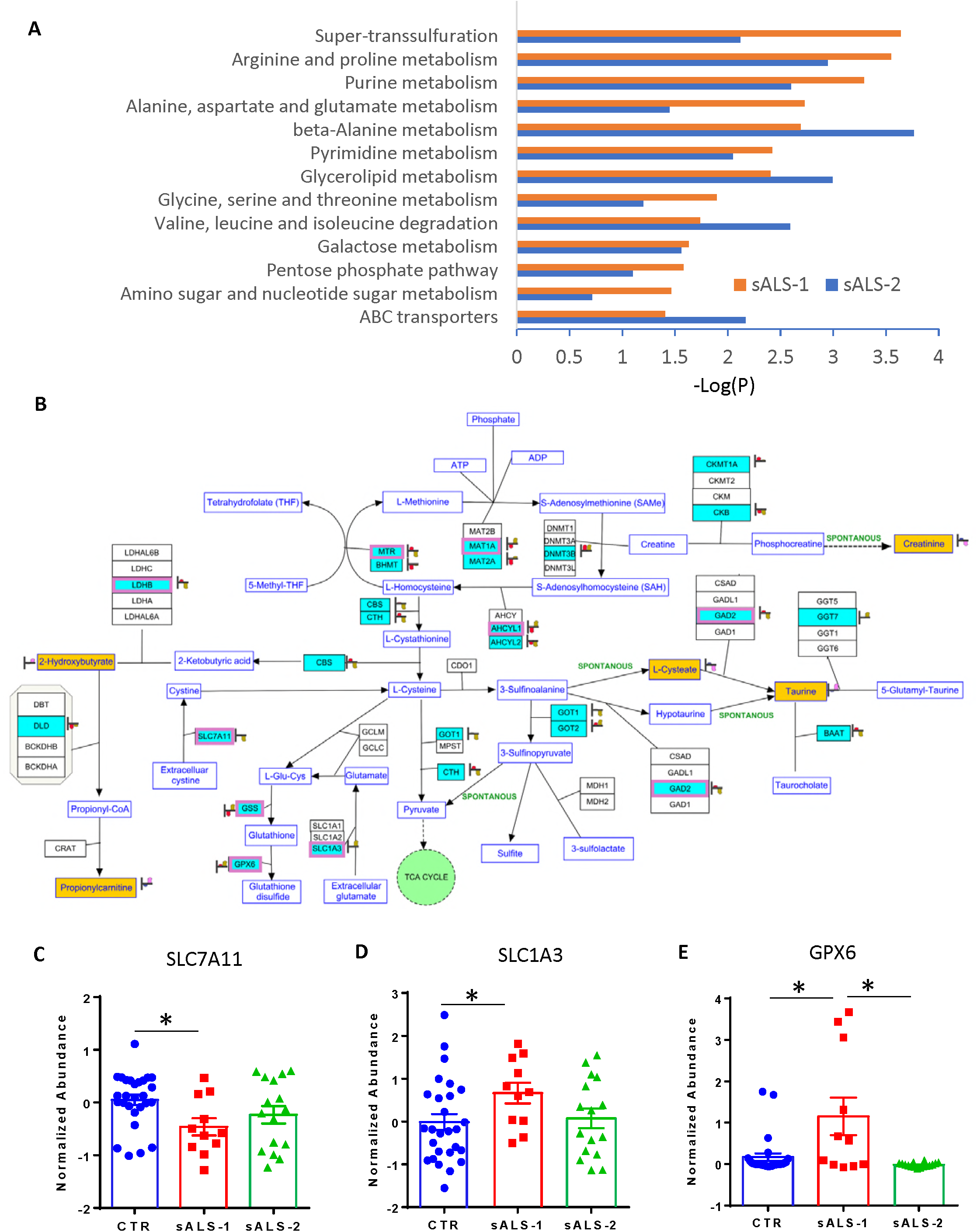
Pathway Integration of transcript and metabolite changes in stratified sALS-1 and non-stratified sALS-2 compared to controls. Panel A: Significant enrichments of integrated multiomic data into metabolic pathways in sALS-1 and sALS-2 compared to control group (P<0.05, Fisher’s exact test). Differentially expressed messenger RNA(mRNA) transcripts and mircroRNA (miRNA) targets were obtained from 11 sALS-1, 16 sALS-2 and 27 age and gender matched controls. Differentially expressed intracellular metabolites were obtained from 18 sALS-1 relative to 18 controls. The metabolic pathways used for mapping were queried by KEGG, Wiki-pathways and Biocyc and imported into Agilent GeneSpring 14.9.1 for pathway analysis. Panel B:Visualization of significant mapping of the differentially expressed mRNA, miRNA targets and metabolites involved in the super-trans-sulfuration pathway in sALS-1 compared to control group (P<0.00024, Fischer’s exact test). Green entities with purple boxes: differentially expressed mRNA (P<0.05); green entities without purple box: differentially expressed miRNA targets (P<0.05); yellow entities: intracellular metabolites. Heat strips next to entities represent normalized abundance in control (left heat strip) and sALS-1 (right heat strip). Panel C-E: Distinctive expression of mRNAs related to cysteine and glutamate transport and glutathione oxidation in sALS-1 compared to sALS-2 and control group. (C): cystine-glutamate antiporter (SLC7A11); (D) glutamate aspartate transporter (SLC1A3); and (E): glutathione peroxidase 6 (GPX6). *, P<0.05 (2-tailed Student t-test).

We next mapped the multiomics data (i.e. differentially expressed mRNA, miRNA targets and metabolites in sALS-1 compared to control) representing the trans-sulfuration pathway to concomitantly visualize transcriptional regulation along with metabolite changes in sALS-1 fibroblasts (Fig 7B). Differentially-expressed mRNA transcripts (entities in green with purple boxes, P<0.05) and differentially expressed microRNA targets (entities in green without purple boxes, P<0.05) were involved not only in the methionine cycle, and homocysteine to cysteine trans-sulfuration, but also in glutathione and cysteine metabolism – both synthesis and oxidation/degradation. We found that sALS-1, but not sALS-2 fibroblast groups showed significant transcriptional changes related to glutathione synthesis and oxidation, compared to the control group (Figs 7C-E).

The cystine/glutamate antiporter (system xc-, SLC7A11) is critical for the generation of GSH by transporting cystine into the cell. System xc-also releases glutamate, which can potentially lead to excitotoxicity. The dual actions of system xc-make it of great interest in ALS with the involvement of both oxidative stress and excitotoxicity. SLC7A11 has previously been shown to be dysregulated in ALS patients and animal models of disease ^22^-^24^. Notably, significant downregulation of SLC7A11 was found in sALS-1, but not sALS-2 fibroblasts, compared to control (Fig 7C). Conversely, SLC1A3, was significantly upregulated in sALS-1, but not in sALS-2, compared to controls (Fig 7D). SLC1A13, also known as the glutamate aspartate antiporter is localized to the plasma membrane as well as the inner mitochondrial membrane where it contributes to the malate-aspartate shuttle by enabling glutamate transport into cells against a concentration gradient. GPX6, an isoform of glutathione peroxidase involved in the oxidation of glutathione to glutathione disulfide, was also found to increase in sALS1, but not sALS-2 fibroblasts, vs. controls (Fig 7E). Increased glutathione oxidation, with an associated decrease in cystine uptake, due to progressive depletion from the culture medium, would predictably result in increased cysteine demand via trans-sulfuration (from homocysteine) for glutathione synthesis. Such an increase in trans-sulfuration may be expected to drive an accumulation of downstream trans-sulfuration pathway metabolites (e.g., taurine, 2-hydroxybutyrate,propionylcarnitine) as observed in sALS-1 fibroblasts. Additionally, upregulation of SLC1A3 might point to an increased malate aspartate shuttle, consistent with the increased glucose flux into the TCA cycle found in sALS1 fibroblasts.

Taken together, sALS-1 case-derived fibroblasts show distinct transcriptional changes involved in cystine and glutamate transport and glutathione synthesis/oxidation, as compared to non-stratified sALS-2 and control fibroblasts.

### sALS patient plasma displays altered trans-sulfuration pathway metabolites

The plasma metabolome of ALS patients has been previously studied in attempt to identify disease biomarkers. We sought to explore if sALS-1 patient plasma exhibit indications of altered metabolism in accord with those identified in their cognate fibroblasts. Plasma samples from the same 18 sALS-1 cases that yielded fibroblasts with aberrant transulfuration/hyper-metabolism phenotype were compared to plasmas from 20 age- and gender-matched controls. Targeted and untargeted metabolite profiling detected >1000 metabolite features, of which 85 were found to be differentially-abundant in the sALS-1 group compared to controls (Fig 8A). PCA analysis revealed that the sALS-1 group cluster separated from the control group, based on relative levels of these 85 metabolites (Fig 8B). Notably, the differences in plasma metabolites in sALS-1 cases included decreased levels of some key amino acids (glutamine, glutamate, methionine, tyrosine) and creatinine, along with increased levels of taurine-related species (i.e., taurine and glutamyl-taurine. bile acids and 2-hydroxybutyrate), purine metabolites, phospholipids, ceramides and fatty acid amides. Two species that were initially unknown and found to be present in the majority of sALS cases were subsequently identified as Riluzole and its primary metabolite, hydroxyriluzole glucuronide. Mapping all identified differentially abundant sALS plasma metabolites to KEGG pathways, revealed the taurine/hypotaurine metabolic pathway to have the highest pathway impact score and the lowest pathway matching p-value (Fig 8C), consistent with the results from fibroblast studies. Notably, increased plasma taurine and 2-hydroxybutyrate (both trans-sulfuration pathway intermediates), along with decreased creatinine (methionine cycle-related methylation product; Fig 1F), exhibited similar patterns of change to observations in fibroblasts, supporting a systemic metabolic perturbation in sALS-1 cases and the potential for effective use of plasma profiling for facile clinical recognition of the sALS-1 metabotype.

**Figure 8.**
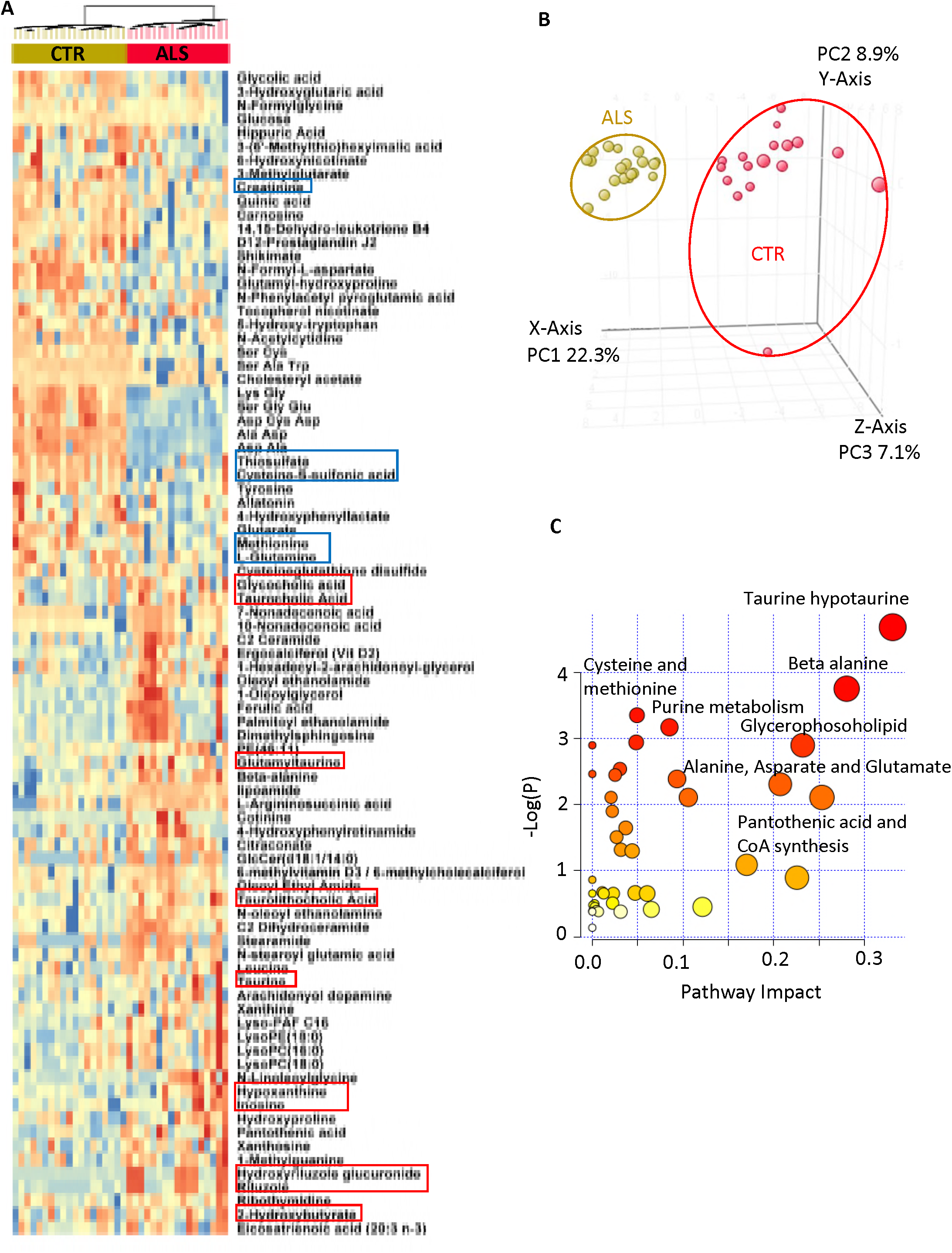
sALS-1 showed distinct plasma metabolite profiles from control and indications of altered taurine and hypo taurine metabolic pathway. Panel A: heatmap of differential plasma metabolites in sALS compared to control. Metabolites related to altered taurine and trans-sulfuration pathway, amino acids, lipid metabolism and purine nucleosides were highlighted in red (upregulated) and blue (downregulated) boxes. sALS drug Riluzole and its drug metabolite hydroxyriluzole glucuronide were also highlighted. Panel B: PCA score plot showing the separation of sALS-1 subgroup from control based on the 88 differentially expressed plasma metabolites. Panel C: Mapping the differentially expressed metabolite into KEGG pathway using Pathway analysis module of MetaboAnalyst 3.0.

## Discussion

In an effort to identify metabolic biomarkers for stratification of sALS patients, we applied untargeted metabolite profiling to dermal fibroblasts isolated from cases. This effort confidently identified a subgroup of sALS fibroblasts (i.e., the sALS-1 group) with a distinctly upregulated trans-sulfuration pathway and related species. Importantly, we could identify this metabotype in both fibroblasts and plasma from cognate sALS-1 cases and found that this subgroup also exhibits perturbed mRNAs for trans-sulfuration/cysteine metabolism compared with control and sALS-2 patients that did not stratify into the sALS-1 group. We further demonstrated that the altered metabolic pathways in the ALS-1 subgroup was associated with increased GSH synthesis and accelerated glucose metabolism. The interplay between increased glucose hypermetabolism and increased antioxidant defense demand, raises the possibility that upregulated antioxidant defenses in sALS-1 cases serves to meet the hypermetabolic demand of this patient subgroup, potentially contributes to disease modulation.

It is known that a subset of ALS patient brains has approximately 10% more resting energy expenditure than healthy individuals^25^-^27^. Notably, hypermetabolism is a feature of more than half of the ALS population and was recently shown to be a deleterious prognostic factor for ALS. Indeed, patients with hypermetabolism have a 20% worse prognosis than those that do not exhibit the hypermetabolism phenotype^28,29^ Until now, hypermetabolism in ALS has mostly been considered in relation to glucose metabolism. Indeed, ^18^F-FDG-PET imaging has been widely applied clinically to assess and discriminate CNS differences in glucose uptake that distinguish hypermetabolism in both familial and sALS patients^25, 30, 31^ Enhanced ^18^F-FDG-PET uptake was found to correlate with cognitive impairment in ALS patients and provides a convenient means to identify altered metabolic activity^30^.

We previously demonstrated that mitochondrial bioenergetics is perturbed in skin fibroblasts from a group of 171 sALS patients, compared with 98 control fibroblast cell lines^5^. Particularly, sALS fibroblasts showed a significant increase in mitochondrial membrane potential (MMP) as well as the ratio between MMP and mitochondrial mass (MMP:MM ratio). Since the sALS patients and control subjects utilized in the present study were part of the same cohort previously studied, we performed meta-analysis of the prior bioenergetics data relative to the sALS-2 subset and control fibroblasts. Based on this analysis, the sALS-1 subgroup,presenting with the enhanced trans-sulfuration metabotype, also exhibited a lower MM, increased MMP, and increased MMP:MM ratio compared to both controls and the sALS-2 subgroup (Fig S8). Interestingly, metabolite profiling and glucose tracing of ribose incorporation into ATP synthesis revealed that the total intracellular ATP pools did not differ among these groups, despite a significantly increased ribose incorporation into ATP (Fig 6). This finding is in agreement with the published reports using a chemiluminescent approach to quantify relative potential differences in ALS vs. control fibroblast levels of ATP^32^. Together, previously reported data suggest that sALS cases require a higher mitochondrial membrane potential as an adaptation to increased ATP demands^5^.

Here, using metabolite profiling and glucose tracing we demonstrate that sALS fibroblasts can be stratified metabolically, based on trans-sulfuration pathway flux and associated accelerated rates of GSH synthesis and glucose metabolism. Moreover, increased intracellular and plasma levels of 2-hydroxybutyrate correlate with increased GSH synthesis and increased taurine abundance in our newly-identified sALS-1 case subgroup. Since limited cystine uptake by fibroblasts appears to accelerate the conversion of homocysteine to cysteine via the trans-sulfuration pathway, fueling production of both GSH and taurine, we speculate that sALS-1 cases arise due to greater demand for anti-oxidant GSH synthesis, owing to a hypermetabolism that produces greater levels of ROS.

We hypothesize that increased glucose metabolism and GSH synthesis are respectively a cause and consequence of oxidant stress in approximately 25% (18/77) of the sALS fibroblast lines that we studied. In accord with this finding, a recent study correlated *in vivo* ALS staging with glucose metabolism patterns and demonstrated that four neuropathological stages of ALS correlate with discriminative regional brain glucose metabolism patterns and with both disease duration and forced vital capacity^33^. Therefore, sALS1 cases are likely to comprise a heterogenous spectrum of glucose hypermetabolism phenotypes, with a distinct metabotype identified herein tuned by cysteine availability to enable antioxidant defenses that evade oxidative stress.

The finding that similar changes in the trans-sulfuration pathway metabolites occur in both fibroblasts and plasma from sALS-1 cases is intriguing and may provide an opportunity for facile clinical stratification of sALS cases based on plasma metabolite profiling. Remarkably, changes in plasma levels 2-hydroxybutyrate,taurine, amino acids and creatinine have been previously reported in ALS patients^34^-^37^. As mentioned above, 2-hydroxybutyrate derives from 2-ketobutyrate, produced from cystathionine in the trans-sulfuration pathway via a NADH-dependent reduction catalyzed by lactate dehydrogenase B (Fig 1F). Elevated levels of plasma 2-hydroxybutyrate and 2-ketobutyrate have previously been shown in several studies involving different cohorts of ALS and ascribed to disease-associated oxidative stress ^34^-^36^. Serum creatinine, a non-enzymatic catabolic product of creatine phosphate that is primarily derived from muscle metabolism at a fairly constant bodily rate (1%-2% of muscle creatine is converted daily to creatinine, followed by transport to the kidney^38^. Decreased plasma creatinine has previously been suggested to be an ALS risk factor^39, 40^

Besides increased plasma taurine, bile acids (e.g., cholic acid) and bile acid conjugates, (e.g.,taurocholic acid), were also found to increase in the sALS-1 subgroup (Fig 9A). Bile acids have been long known to play a neuroprotective role in a diverse spectrum of age-related neurodegenerative disorders^41^. There are two primary bile acids produced by the liver in humans: cholic acid (CA) and chenodeoxycholic acid (CDCA). These primary bile acids can undergo conjugation with glycine or taurine prior to secretion in the bile as glycocholic acid (GCA), taurocholic acid (TCA), glycochenodeoxycholic acid (GCDCA), and taurochenodeoxycholic acid (TCDCA). Therefore, upregulated plasma levels of taurine and taurocholic acid could represent a defense mechanism against heightened oxidative stress.

In summary, we describe the first application of untargeted metabolite profiling and stable isotope tracing as a means to stratify, predict and characterize a subgroup of sALS skin fibroblasts, identifying a distinct metabotype typified by increased trans-sulfuration and glucose hypermetabolism. Because these metabotypic changes were similarly observed in plasma from these sALS patients, we infer that facile plasma profiling can be employed to confidently stratify sALS cases for recognition of the sASL-1 subtype for future to-be-defined targeted anti-oxidant therapies. We hypothesize that the metabolic rewiring that results in upregulation of the trans-sulfuration pathway in these patients arises as an adaptation to limiting cysteine and hence GSH, potentially triggered by hypermetabolism that accelerates GSH oxidation. Knowledge obtained from metabotypic stratification of skin-derived fibroblasts from sALS-1 cases may provide a much-needed first step toward developing precision medicine for personalized therapy of sALS patients.

## Acknowledgements

Funding: NIH R01NS093872 (GM, SG, and LS), R01NS062055 (GM). We thank Dr. Hiroshi Mitsumoto (Columbia University) and the COSMOS initiative for providing the fibroblast and plasma samples utilized in this work.

## Author contributions

DS, BIS, RRC performed metabolite extraction and targeted metabolite analysis, DR and SMF analyzed the transcriptomics data. CK, KB, and HK performed cell culture and the fibroblast bioenergetics analysis. ELC and LC contributed to the data interpretation and concepts presented; QC, GM and SSG conceived the experiments, analyzed the data and wrote the initial manuscript draft for editing by co-authors.

---

**Figure S1.**
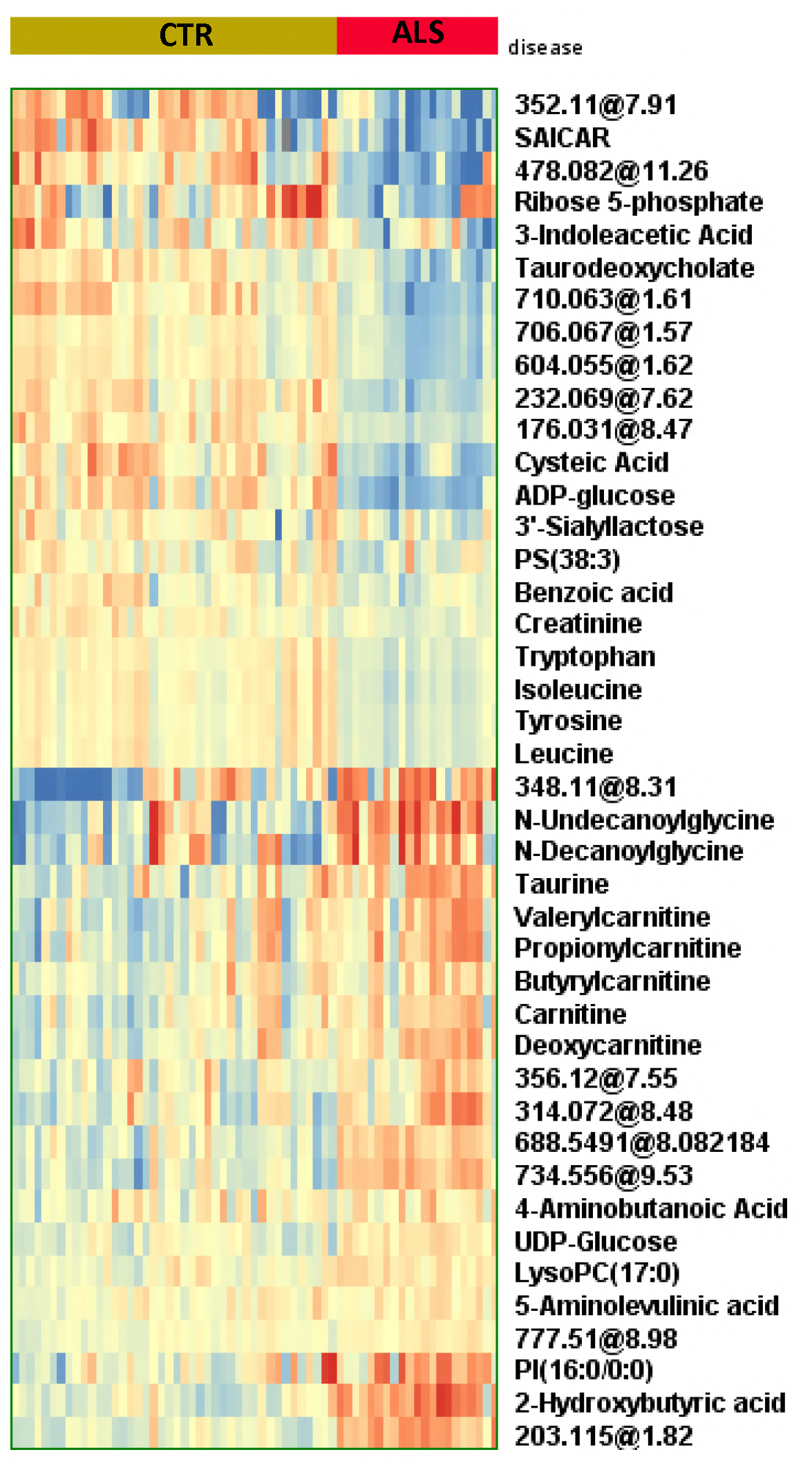
Heatmap depiction of differential metabolites in 18 sALS-1 subgroup compared to 45 control. Unknown metabolites were expressed as accurate mass and retention time. Blue denotes downregulation and red denotes upregulation.

**Figure S2.**
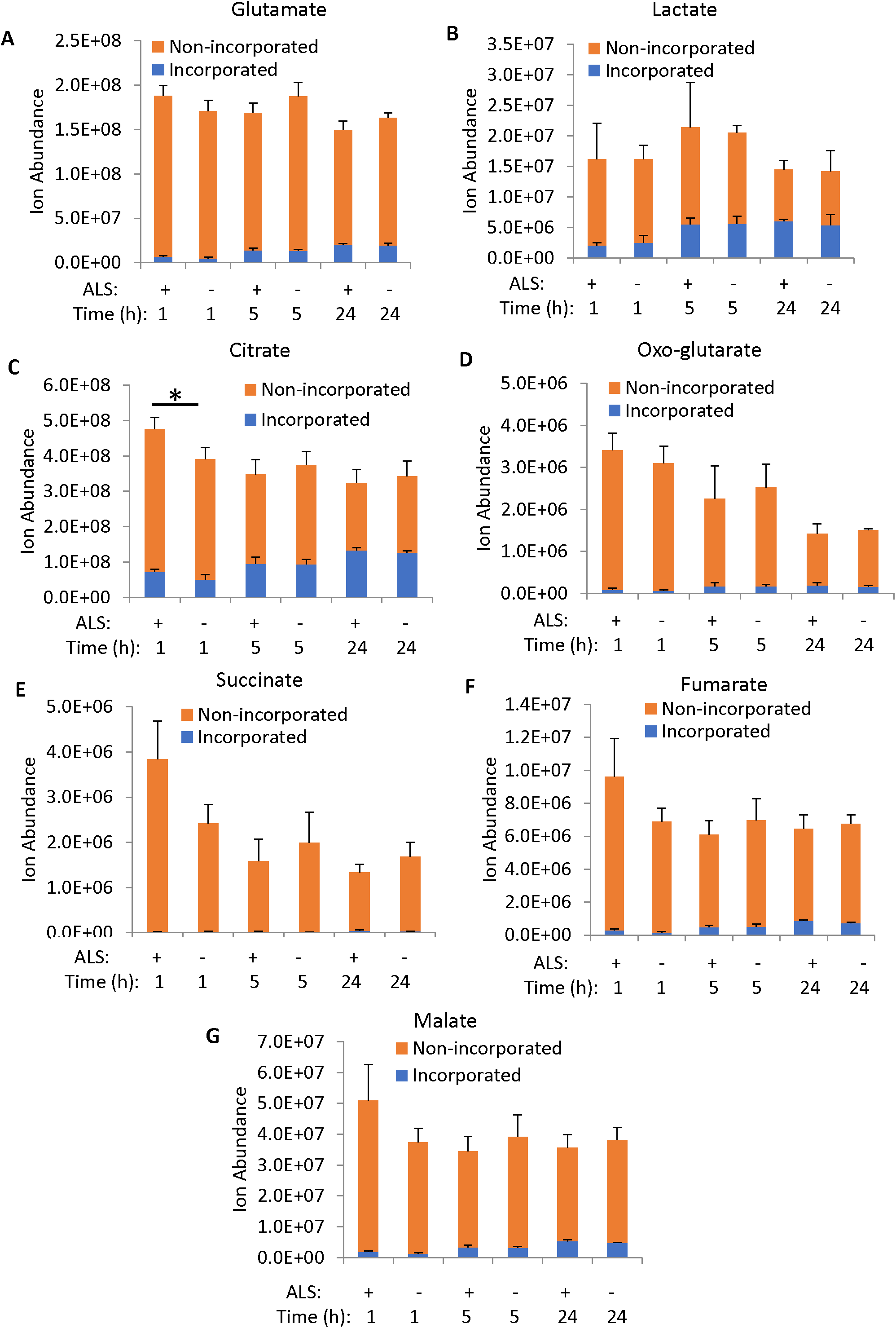
Ion abundance of glutamate, lactate and TCA cycle intermediates after 1h, 5h, and 24h of [U-13C] glucose enrichment. Data was obtained from 3 independent sALS vs. 3 control cell lines and presented as mean ± S.D. P<0.05 (2-tailed Student t-test).

**Figure S3.**
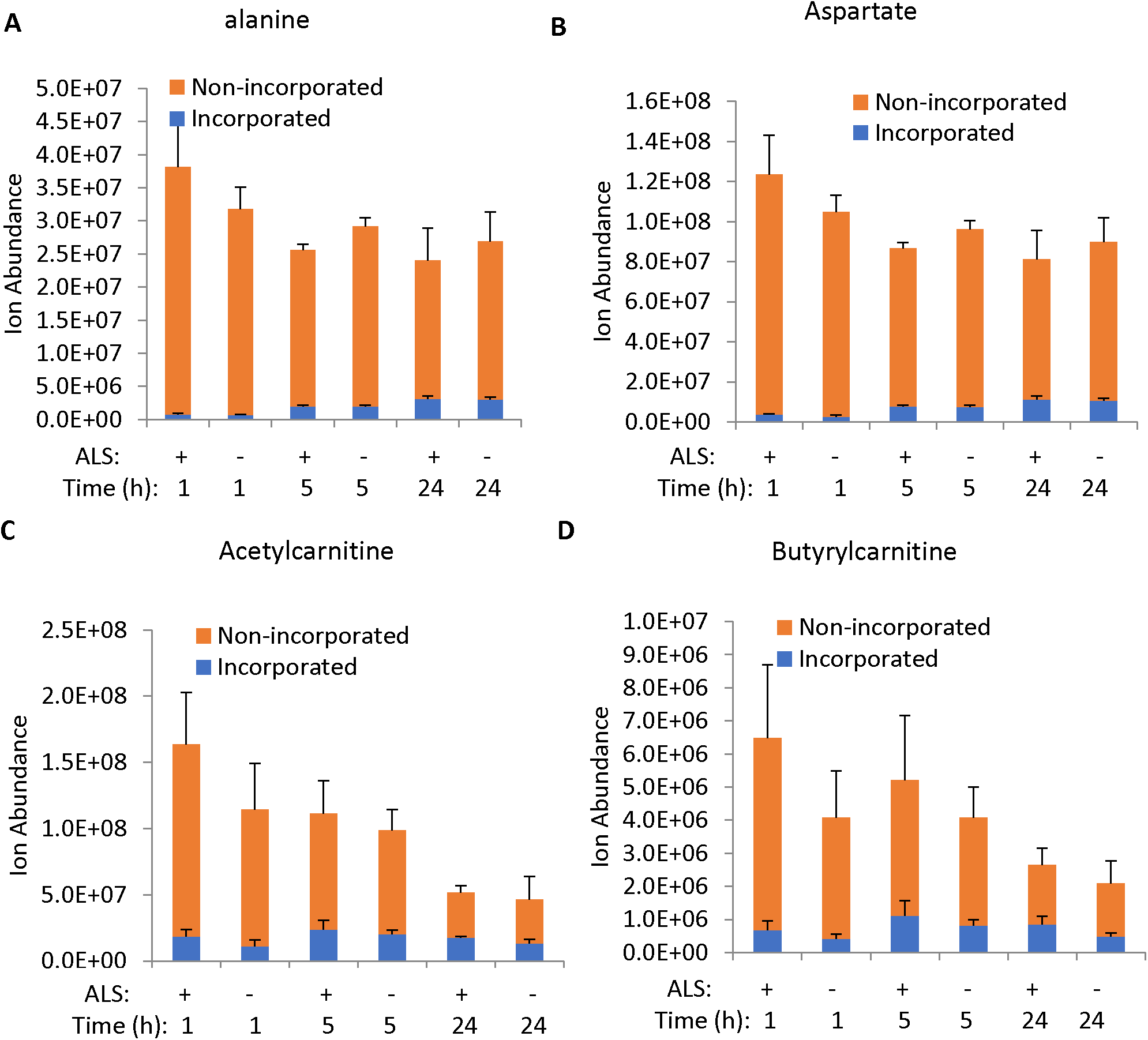
Ion abundance of amino acids and acylcarnitines after 1h, 5h, and 24h of [U-13C] glucose enrichment. Data was obtained from 3 independent sALS vs. 3 control cell lines and presented as mean ± S.D.

**Figure S4.**
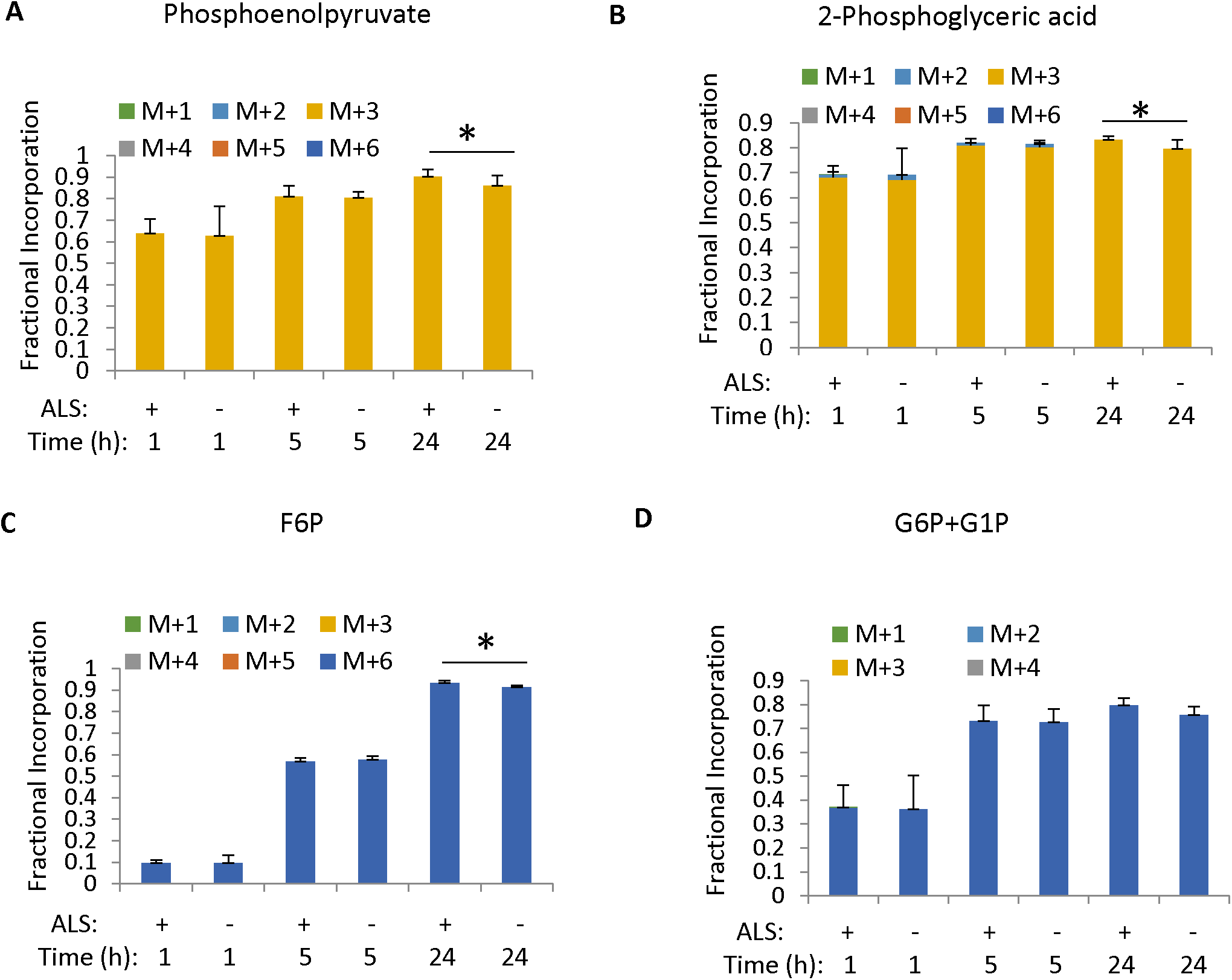
Fractional incorporation of [U-^13^C] glucose into glycolysis intermediates after 1h, 5h and 24h of isotope enrichment. Data was obtained from 3 independent sALS vs. 3 control cell lines and presented as mean ± S.D. *, P<0.05 (2-tailed Student t-test).

**Figure S5.**
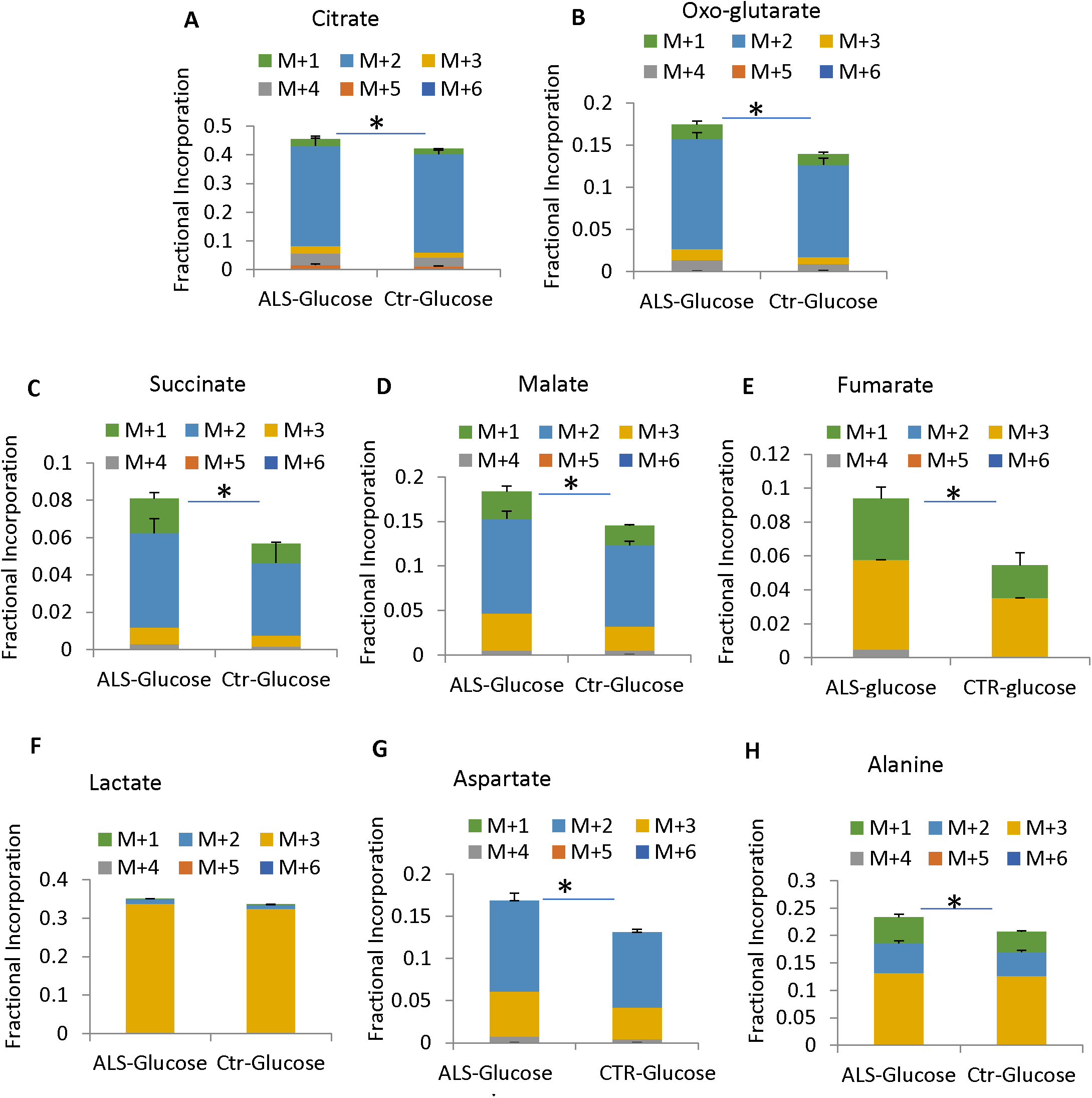
Fractional incorporation of [U-^13^C] glucose into TCA cycle, and amino acids after 48h of isotope enrichment. Data was obtained from 3 independent sALS vs. 3 control cell lines and presented as mean ± S.D.*, P<0.05(2-tailed Student t-test).

**Figure S6.**
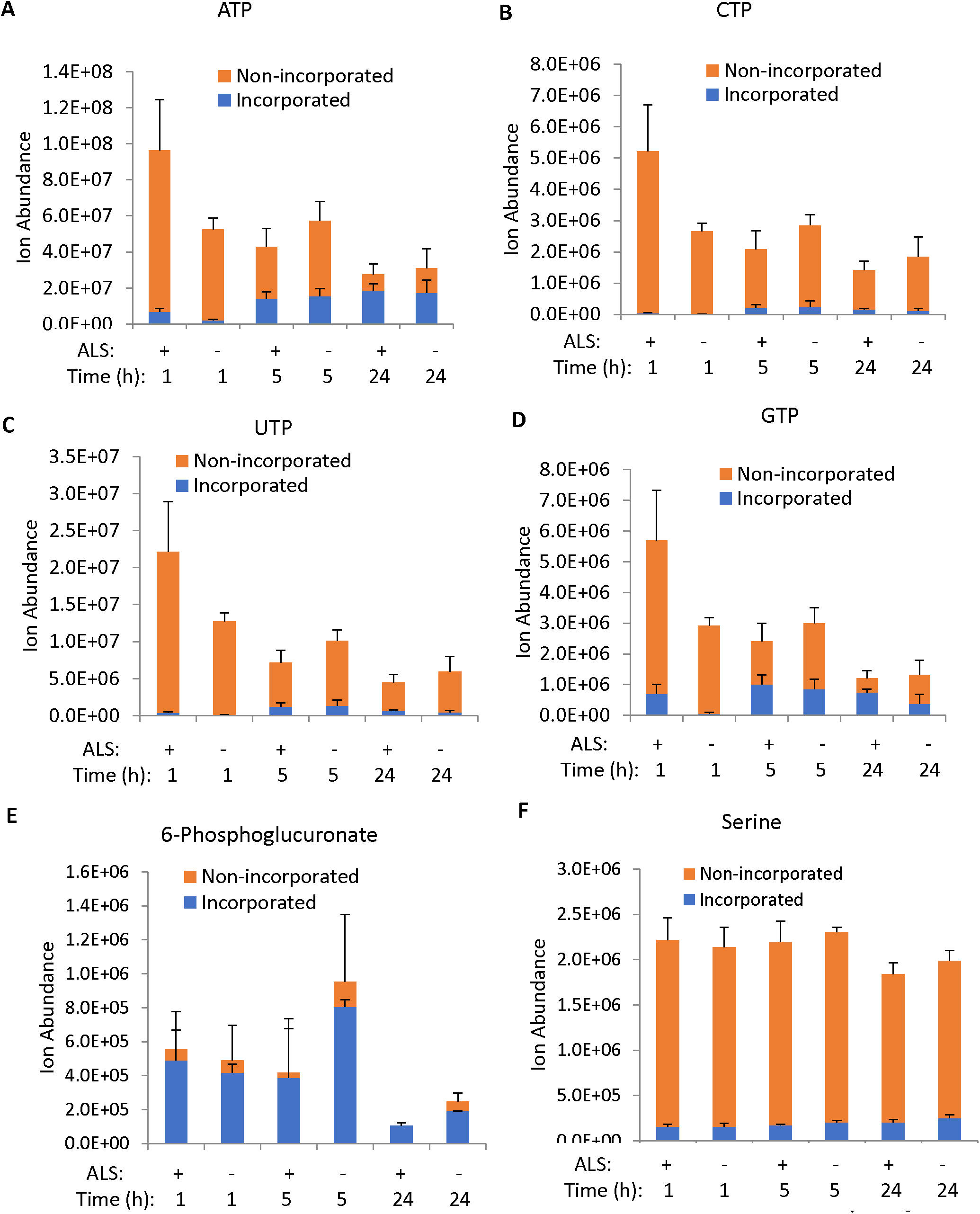
Ion abundance of nucleotide triphosphates after 1h, 5h, and 24h of [U-^13^C] glucose enrichment. Data was obtained from 3 independent sALS vs. 3 control cell lines and presented as mean ± S.D.

**Figure S7.**
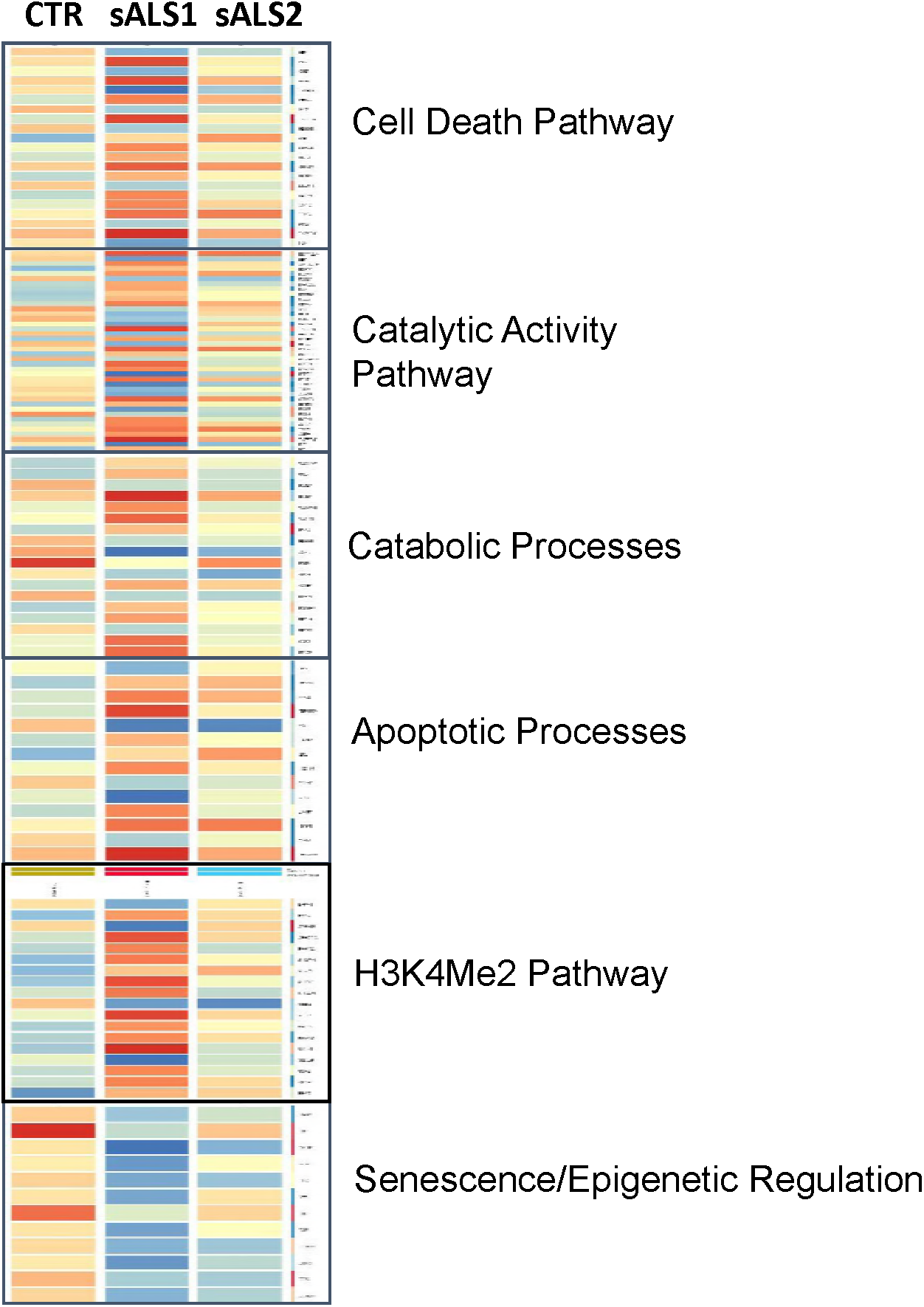
Gene Set Enrichment analysis (GSA) identified significantly different expression of genes involved in the regulation of catabolic processes, apoptosis and epigenetic regulations in sALS-1 compared to sALS-2 and control group.

**Figure S8.**
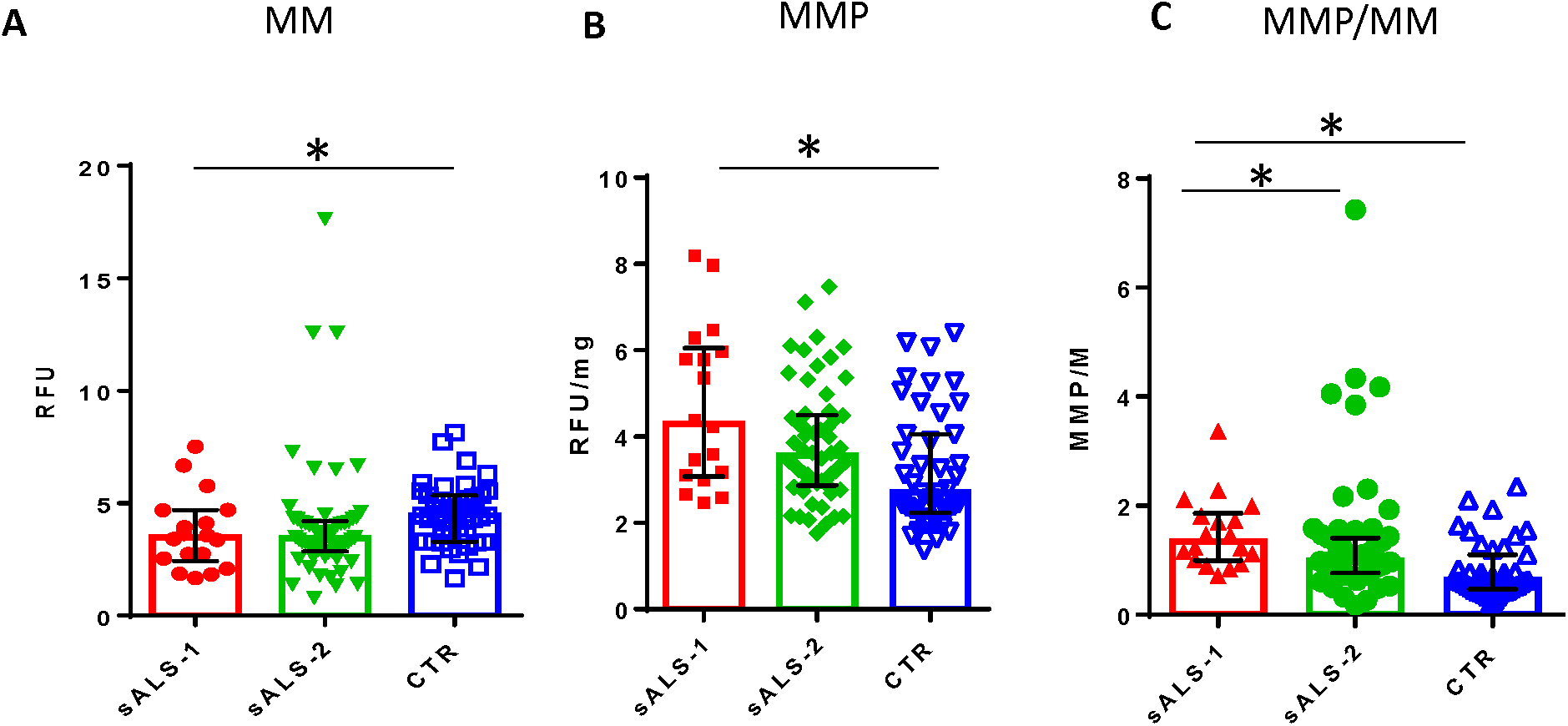
Stratified sALS-1 subgroup showed increased mitochondria membrane potential and mitochondria membrane potential to mass ratio compared to control and non-stratified sALS-2 patients. The numbers of sALS-1, sALS-2 and controls are 18, 58 and 43, respectively. *, P<0.05 (2-tailed Student t-test).

